# The mechanism of RNA capping by SARS-CoV-2

**DOI:** 10.1101/2022.02.07.479471

**Authors:** Gina J. Park, Adam Osinski, Genaro Hernandez, Jennifer L. Eitson, Abir Majumdar, Marco Tonelli, Katie Henzler-Wildman, Krzysztof Pawłowski, Zhe Chen, Yang Li, John W. Schoggins, Vincent S. Tagliabracci

**Affiliations:** Department of Molecular Biology, University of Texas Southwestern Medical Center, Dallas, TX 75390, USA; Department of Microbiology, University of Texas Southwestern Medical Center, Dallas, TX 75390, USA; Department of Biochemistry, University of Wisconsin-Madison, Madison, WI 53706, USA; Department of Biochemistry and Microbiology, Institute of Biology, Warsaw University of Life Sciences, Warsaw 02-776, Poland; Department of Biophysics, University of Texas Southwestern Medical Center, Dallas, TX 75390, USA; Harold C. Simmons Comprehensive Cancer Center, University of Texas Southwestern Medical Center, Dallas, Texas 75390, USA; Hamon Center for Regenerative Science and Medicine, University of Texas Southwestern Medical Center, Dallas, Texas 75390, USA; Howard Hughes Medical Institute, University of Texas Southwestern Medical Center, Dallas, Texas 75390, USA

## Abstract

The SARS-CoV-2 RNA genome contains a 5’-cap that facilitates translation of viral proteins, protection from exonucleases and evasion of the host immune response^1-4^. How this cap is made is not completely understood. Here, we reconstitute the SARS-CoV-2 ^7Me^GpppA_2’-O-Me_-RNA cap using virally encoded non-structural proteins (nsps). We show that the kinase-like NiRAN domain^5^ of nsp12 transfers RNA to the amino terminus of nsp9, forming a covalent RNA-protein intermediate (a process termed RNAylation). Subsequently, the NiRAN domain transfers RNA to GDP, forming the cap core structure GpppA-RNA. The nsp14^6^ and nsp16^7^ methyltransferases then add methyl groups to form functional cap structures. Structural analyses of the replication-transcription complex bound to nsp9 identified key interactions that mediate the capping reaction. Furthermore, we demonstrate in a reverse genetics system^8^ that the N-terminus of nsp9 and the kinase-like active site residues in the NiRAN domain are required for successful SARS-CoV-2 replication. Collectively, our results reveal an unconventional mechanism by which SARS-CoV-2 caps its RNA genome, thus exposing a new target in the development of antivirals to treat COVID-19.

## Main Text

Coronaviruses (CoVs) are a family of positive-sense, single-stranded RNA viruses that cause disease in humans, ranging from mild common colds to more severe respiratory infections^9^. The most topical of these is severe acute respiratory syndrome coronavirus 2 (SARS-CoV-2), the etiological agent of the ongoing COVID-19 pandemic, which to date has resulted in over 5.7-million deaths and almost 400-million cases globally^10^.

The SARS-CoV-2 RNA genome contains two open reading frames (ORF1a and ORF1ab), which are translated by host ribosomes to form two large polyproteins^2^. These polyproteins are subsequently cleaved by viral proteases to form 16 non-structural proteins (nsp1-16), some of which make up the <u>R</u>eplication-<u>T</u>ranscription <u>C</u>omplex (RTC)^2^. At the core of the RTC is the nsp12 <u>R</u>NA-<u>d</u>ependent <u>R</u>NA <u>p</u>olymerase (RdRp), which is the target of several promising antivirals used to treat COVID-19 including remdesivir^11^ and molnupiravir^12^. In addition to the RdRp domain, nsp12 contains an N-terminal <u>N</u>idovirus <u>R</u>dRp-<u>A</u>ssociated <u>N</u>ucleotidyltransferase (NiRAN) domain (**Fig. 1a**)^5^. The NiRAN domain shares sequence and structural similarity with the pseudokinase selenoprotein-O (SelO), which transfers AMP from ATP to protein substrates (a process termed AMPylation)^13-15^. Notably, the active site kinase-like residues of the NiRAN domain are highly conserved in *Nidovirales* (**Extended Data Fig. 1**) and are required for equine arteritis virus (EAV) and SARS-CoV-1 replication in cell culture^5^. Several hypotheses for the function of the NiRAN domain have been proposed, including roles in protein-primed RNA synthesis, RNA ligation, and mRNA capping^5,16^.

**Figure 1.**
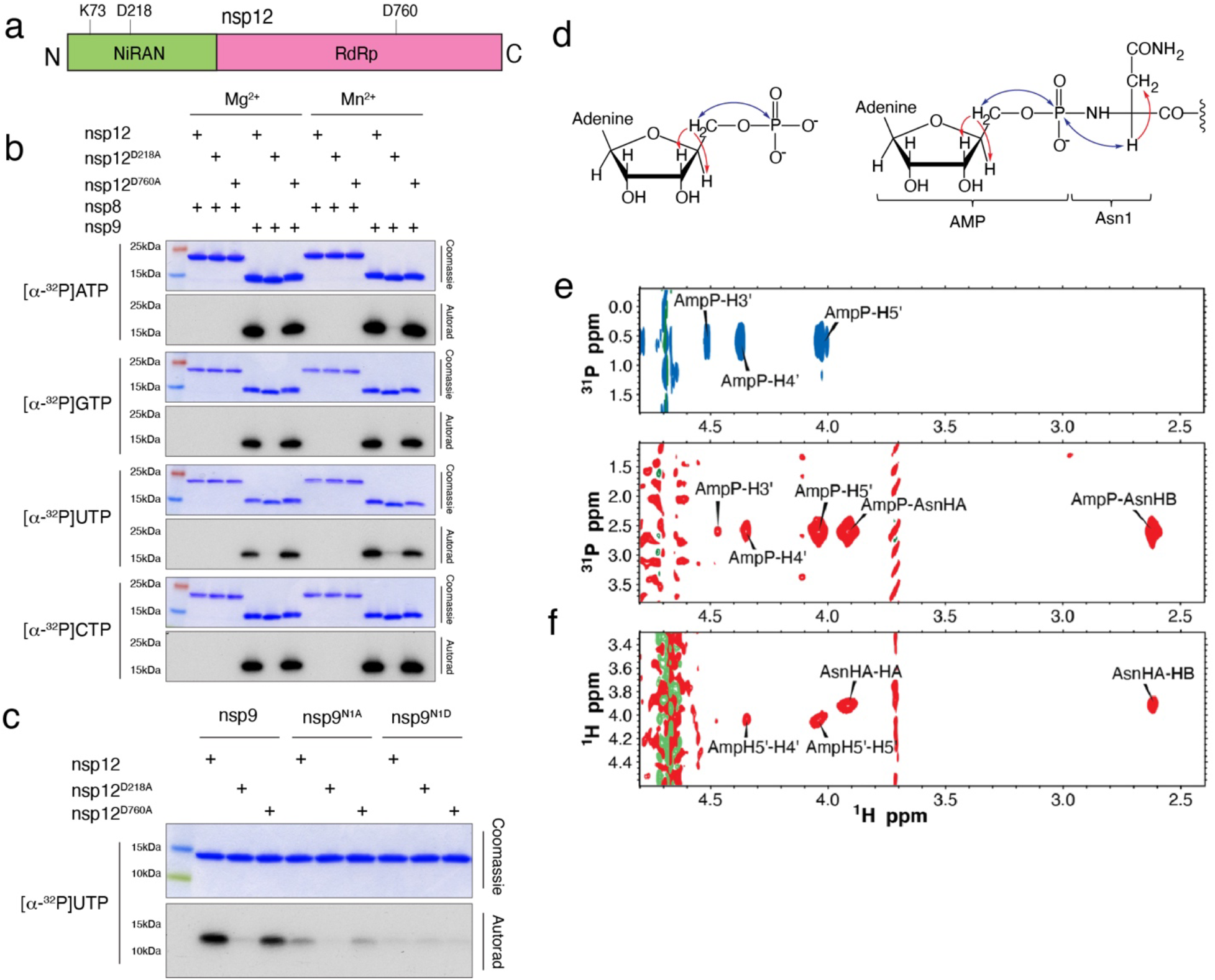
The NiRAN domain NMPylates the N-terminus of nsp9. **a.** Domain architecture of nsp12 depicting the SelO-like NiRAN domain (*green*) and the RNA-dependent RNA polymerase domain (RdRp; *magenta),* annotated with the predicted catalytic residues. **b.** Incorporation of α-^32^P from [α-^32^P]ATP, GTP, UTP, or CTP into nsp8 or nsp9 by WT nsp12, the NiRAN mutant (D218A), or the polymerase mutant (D760A). Reactions were performed in the presence of Mg^2+^ or Mn^2+^ and the products were resolved by SDS-PAGE and visualized by Coomassie staining (*top*) and autoradiography (*bottom).* **c.** Incorporation of α-^32^P from [α-^32^P]UTP into nsp9 or the indicated mutants by the NiRAN domain. Reaction products were analysed as in **b**. **d.** Structure of AMP (*left*) and AMP-nsp9 (*right*) with arrows to indicate the magnetization transfer steps that result in the peaks observed in the 2D NMR spectra. The blue arrows indicate the transfer steps that yield the peaks in the HSQC spectra, while the red arrows show the additional magnetization transfer during the TOCSY that result in the additional peaks found in the HSQC-TOCSY spectra. **e.** 2D 1H,31P-HSQC-TOCSY spectra of AMP (*top, blue*) and AMP-nsp9 (*bottom, red*). **f.** 2D 1H,1H-HSQC-TOCSY spectra of AMP-nsp9. Results shown in **b** and **c** are representative of at least 3 independent experiments.

The CoV RNA genome, like eukaryotic mRNAs, contains a methylated guanosine linked to the first nucleotide of the RNA via a reverse 5’ to 5’ triphosphate linkage (**Extended Data Fig. 2**)^1,4^. This 5’ cap is important for RNA stability, initiation of mRNA translation, and protection from exonucleases^17^. Methylation of the ribose 2’-OH position of the first nucleotide completes the cap and protects the RNA from the host immune system^18,19^. Thus, formation of the RNA cap is crucial for successful replication and transcription of the viral genome.

All eukaryotes share a conserved co-transcriptional capping mechanism (**Extended Data Fig. 2**) involving: **1)** an RNA triphosphatase (RTPase), which removes the g-phosphate from the nascent 5’-triphosphorylated RNA (5’-pppRNA) to yield a 5’-diphosphorylated RNA (5’-ppRNA); **2)** a guanylyltransferase (GTase), which transfers GMP from GTP to 5’-ppRNA to form the cap core structure GpppN-RNA; **3)** a (guanine-N7)-methyltransferase (N7-MTase), which methylates the cap guanine at the N7 position; and **4)** a (nucleoside-2’-*O*)-methyltransferase (2’-O-MTase), which methylates the ribose-2’-OH position on the first nucleotide of the RNA. In CoVs, the nsp13, nsp14, and nsp16 proteins have RTPase^20^, N7-MTase^6^, and 2’-O-MTase^7^ activities, respectively. Thus, it was presupposed that the CoV capping mechanism occurs in a similar fashion to the eukaryotic capping pathway, with the NiRAN domain functioning as the GTase^3,5,21^. However, evidence to support this claim has been lacking.

In this study, we discover that the NiRAN domain transfers monophosphorylated RNA (5’-pRNA) from 5’-pppRNA to the N-terminus of nsp9 as an intermediate step in cap synthesis. The NiRAN domain then transfers 5’-pRNA from RNAylated nsp9 to GDP to form the cap core structure GpppA-RNA. We then reconstitute cap-0 and cap-1 structures using the nsp14 and nsp16 methyltransferases. Furthermore, we present a cryo-EM structure of the SARS-CoV-2 RTC with the native N-terminus of nsp9 bound in the NiRAN active site. Finally, we demonstrate in a reverse genetics system that the N-terminus of nsp9 and the kinase-like active site residues in the NiRAN domain are required for SARS-CoV-2 replication.

### The NiRAN domain NMPylates the N-terminus of nsp9

The NiRAN domain has been shown to transfer nucleotide monophosphates (NMPs) from nucleotide triphosphates (NTPs) (referred to as NMPylation) to protein substrates, including nsp9^16^ and the nsp12 co-factors, nsp7^22^ and nsp8^23^. We observed NiRAN-dependent NMPylation of native nsp9, but not native nsp7 or nsp8 (**Fig. 1b, Extended Data Fig. 3**). Quantification of ^32^P incorporation and intact mass analyses suggests stoichiometric incorporation of NMPs into nsp9 (**Extended Data Fig. 3g-j**). Mutation of nsp9 Asn1 to Ala or Asp reduced NMPylation of nsp9 (**Fig. 1c, Extended Data Fig. 4a**), consistent with previous work that suggested NMPylation occurs on the backbone nitrogen of nsp9 Asn1 ^16^. To provide direct evidence that the amino terminus of nsp9 is NMPylated by the NiRAN domain, we performed nuclear magnetic resonance (NMR) spectroscopy of AMPylated nsp9. The 2D ^1^H,^31^P HSQC and 2D HSQC-TOCSY spectra confirm that the AMP is attached to the nitrogen backbone atom of Asn1 via a phosphoramidate linkage (**Fig. 1d-f, Extended Data 4b, c**).

### The NiRAN domain RNAylates nsp9

Given the ability of the NiRAN domain to transfer NMPs to nsp9 using NTPs as substrates, we wondered whether the NiRAN domain could also utilize 5’-pppRNA in a similar fashion (**Fig. 2a**). We synthesized a 5’-pppRNA 10-mer corresponding to the first 10 bases in the leader sequence (LS10) of the SARS-CoV-2 genome (hereafter referred to as 5’-pppRNA^LS10^) (**Extended Data Table 1).** We incubated 5’-pppRNA^LS10^ with nsp9 and nsp12 and analysed the reaction products by SDS-PAGE. Remarkably, we observed an electrophoretic mobility shift in nsp9 that was time-dependent, sensitive to RNAse A treatment and required an active NiRAN domain, but not an active RdRp domain (**Fig. 2b**). Intact mass analyses of the reaction products confirmed the incorporation of monophosphorylated RNA^LS10^ (5’-pRNA^LS10^) into nsp9 (**Fig. 2c**). The reaction was dependent on Mn^2+^ (**Extended Data Fig. 5a**) and required a triphosphate at the 5’-end of the RNA (**Extended Data Fig. 5b)**. Substituting Ala for Asn1 reduced the incorporation of RNA^LS10^ into nsp9 (**Fig. 2d**). We also observed NiRAN-dependent RNAylation of nsp9 using LS RNAs ranging from 2 to 20 nucleotides (**Fig. 2e**). Mutation of the first A to any other nucleotide markedly reduced RNAylation (**Fig. 2f**). Thus, the NiRAN domain RNAylates the N-terminus of nsp9 in a substrate-selective manner.

**Figure 2.**
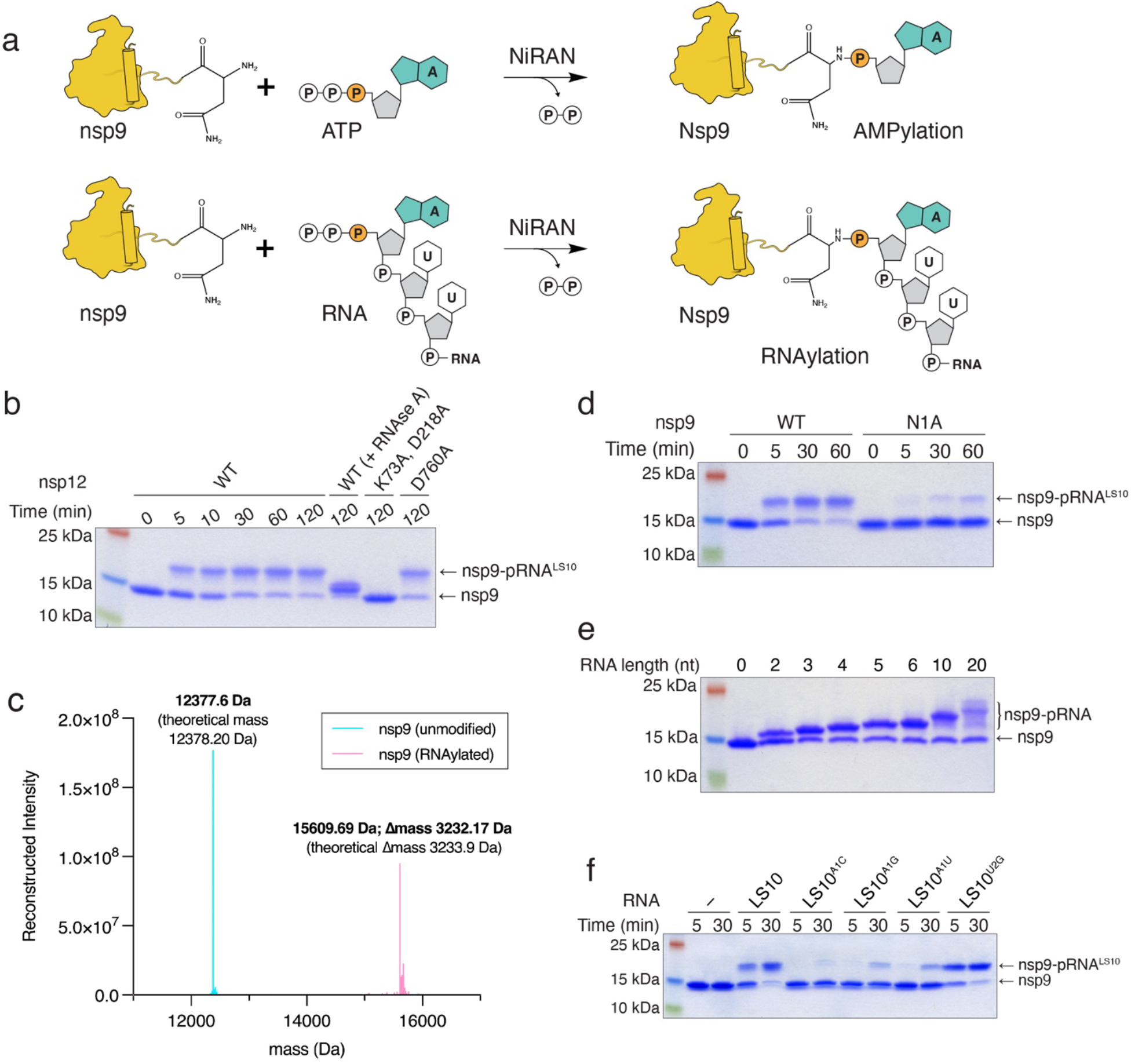
The NiRAN domain RNAylates nsp9. **a.** Schematic depicting the nsp9 AMPylation reaction (*top*) and the proposed nsp9 RNAylation reaction (*bottom*). **b.** Time-dependent incorporation of RNA into nsp9 by WT nsp12, the NiRAN mutant (K73A, D218A), or the polymerase mutant (D760A). Reaction products were analysed by SDS-PAGE and Coomassie staining. Samples were also treated with RNAse A. **c.** Intact mass LC/MS spectra (overlayed) of unmodified nsp9 (*cyan*) or nsp9 after incubation with 5’-pppRNA^LS10^ and WT nsp12 (*pink*). The theoretical and observed masses are shown in the insets. The △mass of 3233.17 Da corresponds to monophosphorylated RNA^LS10^ (5’-pRNA^LS10^). **d.** Time-dependent incorporation of 5’-pRNA^LS10^ into nsp9 or the nsp9 N1A mutant. Reaction products were analysed as in **b**. **e.** Incorporation of different lengths of 5’-pppRNAs into nsp9 by the NiRAN domain. Reaction products were analysed as in **b**. **f.** Time-dependent incorporation of RNAs with substitutions in the first and second base into nsp9 by the NiRAN domain. Reaction products were analysed as in **b**. Results shown are representative of at least 2 independent experiments.

### The NiRAN domain transfers 5’-pRNA from nsp9 to GDP forming the cap core structure GpppA-RNA

Negative-sense RNA viruses of the order *Mononegavirales,* including vesicular stomatitis virus (VSV), have an unconventional capping mechanism in which a polyribonucleotidyltransferase (PRNTase) transfers 5’-pRNA from 5’-pppRNA to GDP via a covalent enzyme-RNA intermediate (**Extended Data Fig. 6a**)^24,25^. Because the NiRAN domain transfers 5’-pRNA to nsp9, we hypothesized that this protein-RNA species may be an intermediate in a similar reaction mechanism to that of the VSV system. To test this hypothesis, we purified the nsp9-pRNA^LS10^ species by ion exchange and gel filtration chromatography and incubated it with GDP in the presence of nsp12. Treatment with GDP deRNAylated nsp9 in a NiRAN-dependent manner, as judged by the nsp9 electrophoretic mobility on SDS-PAGE (**Fig. 3a**) and its molecular weight based on intact mass analysis (**Fig. 3b**). The reaction was time-dependent, (**Fig 3c**), preferred Mg^2+^ over Mn^2+^ (**Extended Data Fig. 6b**) and was specific for GDP—and to some extent GTP—but not the other nucleotides tested (**Fig. 3d**). Interestingly, although inorganic pyrophosphate (PPi) was able to deAMPylate nsp9-AMP, it was unable to deRNAylate nsp9-pRNA^LS10^ (**Fig. 3e**). (See Discussion)

**Figure 3.**
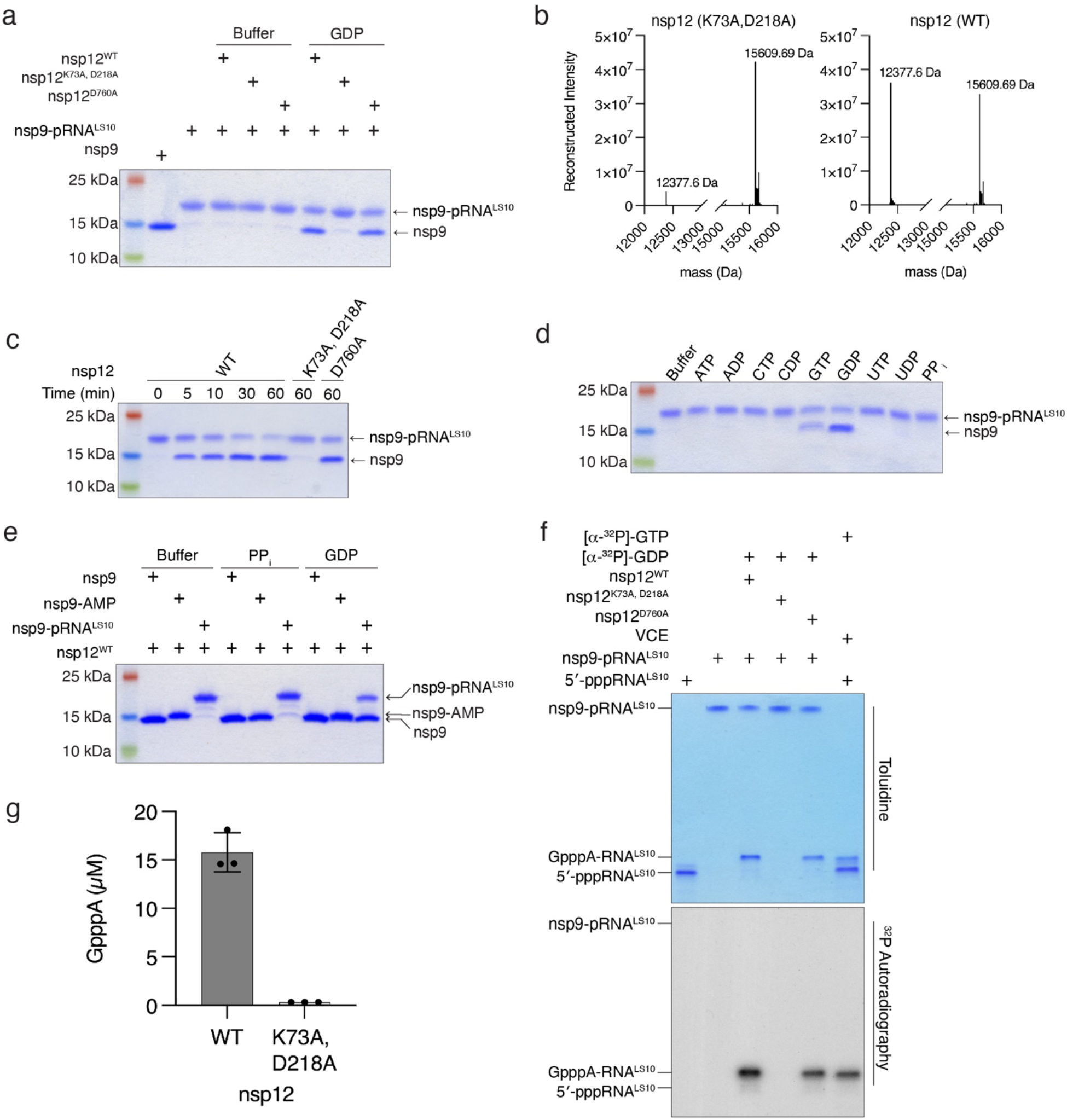
The NiRAN domain catalyses the transfer of 5’-pRNA from nsp9 to GDP to form the cap core structure GpppA-RNA. **a.** DeRNAylation of the covalent nsp9-RNA^LS10^ species by WT nsp12, the NiRAN mutant (K73A, D218A), or the polymerase mutant (D760A) when incubated with buffer or GDP. Reaction products were analysed as in **Fig. 2b**. **b.** Intact mass LC/MS spectra of nsp9-pRNA^LS10^ after incubation with GDP and WT nsp12 (*right*) or the NiRAN mutant (*left*). The theoretical mass of nsp9 is 12378.2 Da and the theoretical mass of nsp9-pRNA^LS10^ is 15611.5 Da. **c.** Time-dependent deRNAylation of nsp9-pRNA^LS10^ by WT nsp12, the NiRAN mutant (K73A, D218A), or the polymerase mutant (D760A). Reaction products were analysed as in **Fig. 2b**. **d.** DeRNAylation of nsp9-pRNA^LS10^ by nsp12 in the presence of different NTPs, NDPs or PPi. Reaction products were analysed as in **Fig. 2b**. **e.** NiRAN-dependent deAMPylation or deRNAylation of nsp9 in the presence of PPi or GDP. Reaction products were analysed as in **Fig. 2b**. **f.** Incorporation of α-^32^P from [α-^32^P]GDP into nsp9-pRNA^LS10^ by WT nsp12, the NiRAN mutant (K73A, D218A), or the polymerase mutant (D760A). Vaccinia capping enzyme (VCE) was used as a control but incubated with [α-^32^P]GTP. Reaction products were resolved by Urea-PAGE and visualized by toluidine blue O staining (upper) and autoradiography (lower). **g.** HPLC/MS quantification of GpppA formed during the NiRAN-catalysed deRNAylation of nsp9-pRNA^LS10^. Reaction products were digested with nuclease P1 prior to HPLC analysis. Reactions were performed in triplicate and error bars represent the standard deviation. Results shown are representative of at least 2 independent experiments.

We used Urea-PAGE to analyse the fate of the RNA^LS10^ during the deRNAylation reaction. Treatment of nsp9-pRNA^LS10^ with nsp12 and [α-^32^P]GDP resulted in a [^32^P]-labelled RNA species that migrated similarly to GpppA-RNA^LS10^ produced by the Vaccinia capping enzyme (**Fig. 3f**). The reaction was dependent on a functional NiRAN domain but not an active RdRp domain. To confirm the presence of a GpppA-RNA cap, we digested the RNA produced from the nsp12 reaction with P1 nuclease and detected GpppA by high performance liquid chromatography/mass spectrometry (HPLC/MS) analysis (**Fig. 3g**). Thus, the NiRAN domain is a GDP polyribonucleotidyltransferase (GDP-PRNTase) that mediates the transfer of 5’-pRNA from nsp9 to GDP.

In our attempts to generate GpppA-RNA^LS10^ in a “one pot” reaction, we found that GDP inhibited the RNAylation reaction (**Extended Data Fig. 6c**). However, the formation of GpppA-RNA^LS10^ could be generated in one pot provided that the RNAylation occurs prior to the addition of GDP (**Extended Data Fig. 6c, d**).

### Nsp14 and nsp16 catalyse the formation of the cap-0 and cap-1 structures

The SARS-CoV-2 genome encodes an N7-MTase domain within nsp14^6^ and a 2’-O-MTase in nsp16, the latter of which requires nsp10 for activity^7^. Nsp14 and the nsp10/16 complex use S-adenosyl methionine (SAM) as the methyl donor. To test whether NiRAN-synthesized GpppA-RNA^LS10^ can be methylated, we incubated ^32^P-labelled GpppA-RNA^LS10^ with nsp14 and/or the nsp10/16 complex in the presence of SAM and separated the reaction products by Urea-PAGE (**Fig. 4a**). We extracted RNA from the reaction, treated it with P1 nuclease and CIP, and then analysed the products by thin layer chromatography (TLC) (**Fig. 4b**). As expected, the NiRAN-synthesized cap migrated similarly to the GpppA standard and the products from the Vaccinia capping enzyme reaction (compare lanes 1 and 4). Likewise, reactions that included SAM and nsp14 migrated similarly to the ^7Me^GpppA standard and to the products from the Vaccinia capping enzyme reaction following the addition of SAM (compare lanes 2 and 6). Furthermore, treatment of ^7Me^GpppA-RNA^LS10^, but not unmethylated GpppA-RNA^LS10^, with nsp10/16 produced the ^7Me^GpppA_2’-O-Me_-RNA cap-1 structure (compare lanes 3, 8 and 9). In parallel experiments, we incubated NiRAN-synthesized GpppA-RNA^LS10^ with nsp14 and/or the nsp10/16 complex in the presence of [^14^C]-labelled SAM (^14^C on the donor methyl group) and separated the reaction products by Urea-PAGE. As expected, nsp14 and the nsp10/16 complex incorporated ^14^C into GpppA-RNA^LS10^ to form the cap-0 and cap-1 structures, respectively (**Fig 4c**). Thus, the SARS-CoV-2 ^7Me^GpppA_2’-O-Me_-RNA capping mechanism can be reconstituted in vitro using virally encoded proteins.

**Figure 4.**
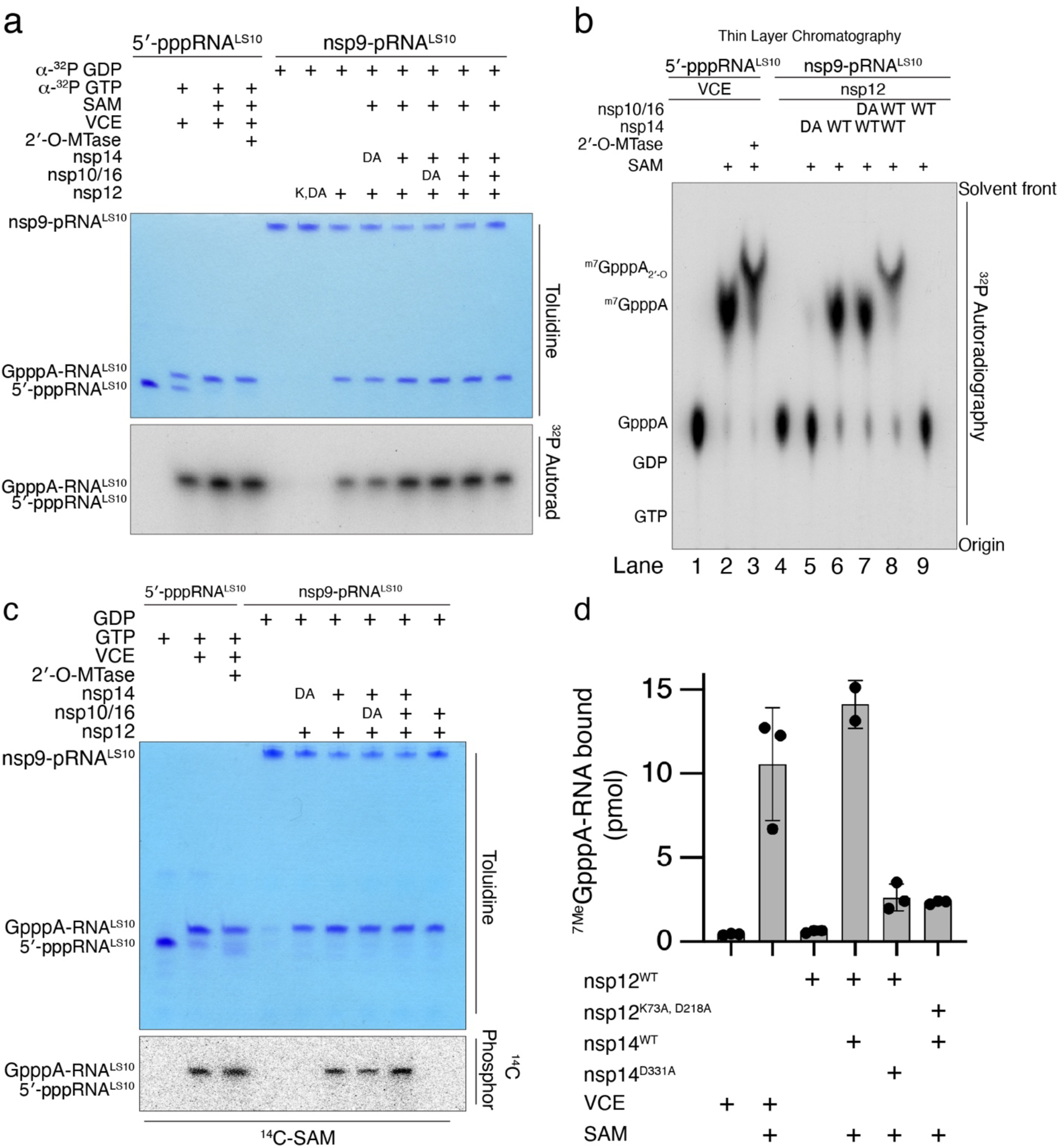
Nsp14 and nsp16 catalyse the formation of the cap-0 and cap-1 structures. **a.** Incorporation of α-^32^P from [α-^32^P]GDP into nsp9-pRNA^LS10^ by WT nsp12, or the NiRAN K73A, D218A mutant (K, DA). Reactions were subsequently incubated with SAM, nsp14 (or the D331A mutant; DA), and nsp10/16 (or the D130A nsp16 mutant; DA). The Vaccinia capping enzyme (VCE) was used as a positive control but incubated with [α-^32^P]GTP and the Vaccinia 2’-O-MTase. Reaction products were analysed as in **Fig 3f**. **b.** Thin layer chromatograms depicting the reaction products from **Fig. 4a** following extraction from the Urea PAGE gel and treatment with PI nuclease and CIP. Location of the cold standards (left) was visualized by UV fluorescence and the ^32^P by autoradiography. **c.** Incorporation of ^14^C from [methyl-^14^C]SAM into GpppA-RNA^LS10^ by nsp14 (or the D331A mutant; DA), and nsp10/16 (or the D130A nsp16 mutant; DA). The VCE and the Vaccinia 2’-O-MTase were used as positive controls. Reaction products were analysed as in **Fig 3f.** The ^14^C signal was detected by phophorimaging. **d.** Pull-down assays depicting the binding of [^32^P]^7Me^GpppA-RNA^LS10^ to GST-eIF4E. [^32^P]^7Me^GpppA-RNA^LS10^ was produced using SARS-CoV-2 virally encoded proteins or the VCE. Radioactivity in GST pull-downs was quantified by scintillation counting. Results represent three independent experiments. Error bars represent the standard deviation (SD). Results shown are representative of at least 2 independent experiments.

Efficient translation of mRNAs is dependent on eIF4E binding to the ^7Me^GpppA-RNA cap^26^. To test whether the SARS-CoV-2 RNA cap is functional, we incubated [^32^P]-labelled ^7Me^GpppA-RNA^LS10^ with GST-tagged eIF4E. We observed [^32^P]-labelled RNA in GST pulldowns of [^32^P]^7Me^GpppA-RNA but not the unmethylated derivative (**Fig. 4d**). Thus, the ^7Me^GpppA-RNA cap generated by SARS-CoV-2 encoded proteins is a substrate for eIF4E in vitro, suggesting that the cap is functional.

### Structural insights into RNA capping by the NiRAN domain

We determined a cryo-EM structure of the nsp7/8/9/12 complex and observed a nsp9 monomer bound in the NiRAN active site (**Fig. 5a, Extended Data Fig. 7-9, Extended Data Table 2**). The native N-terminus of nsp9 occupies a similar position to previously reported structures using a non-native N-terminus of nsp9 (**Fig. 5b, c**)^21^. Our cryo-EM analysis was hindered by the preferred orientation of the complex and sample heterogeneity, yielding final maps with high levels of anisotropy, with distal portions of nsp9 missing, and weak density for the N-lobe of the NiRAN domain (**Extended Data Fig. 7, 8**). Therefore, we used our model and the complex structure by Yan et al.^21^ (PDBID: 7CYQ) to study the structural basis of NiRAN-mediated RNA capping.

**Figure 5.**
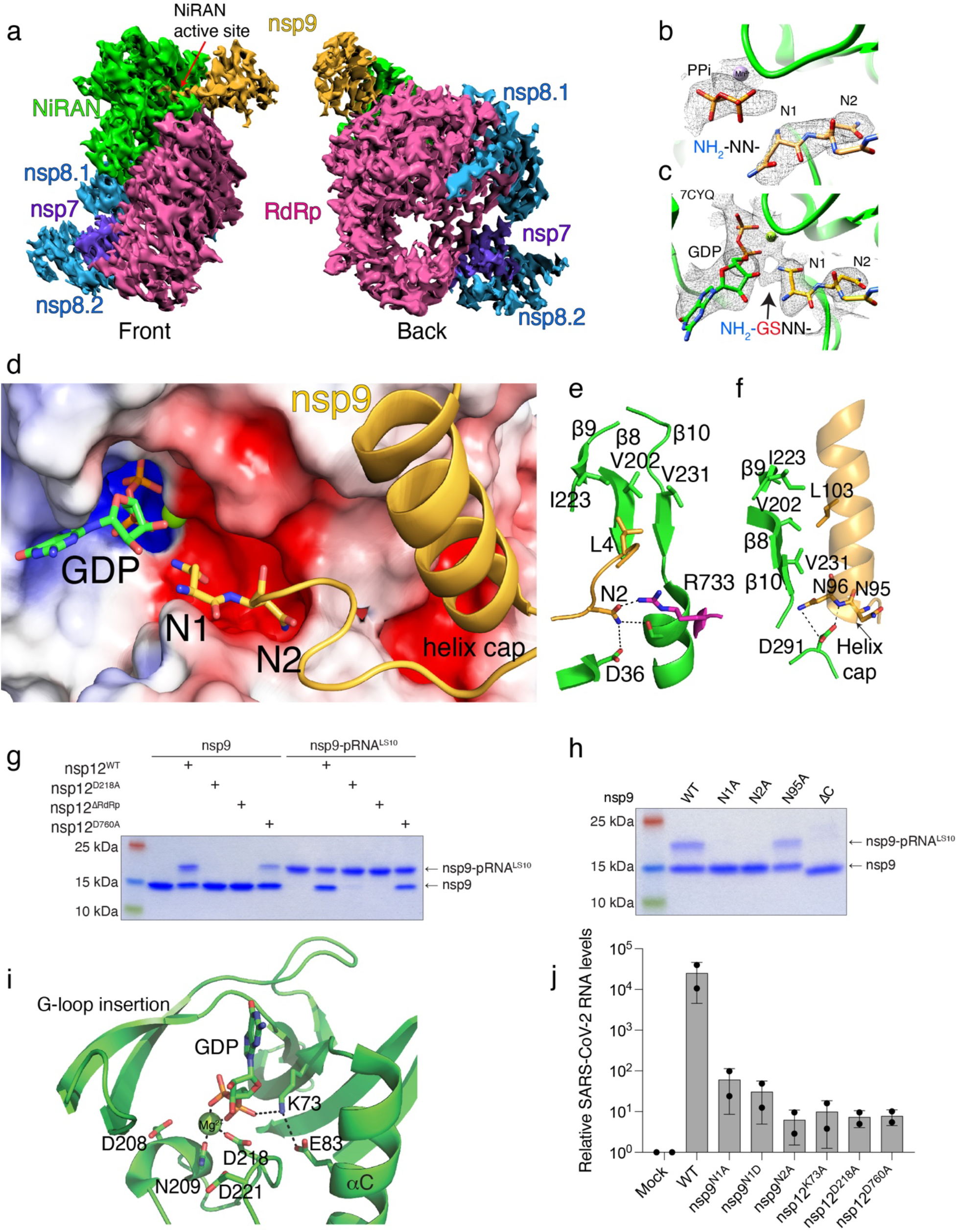
Structural and genetic insights into RNA capping by the kinase domain. **a.** Front and back views of nsp12/7/8/9 cryo-EM maps, with respect to the NiRAN domain. The NiRAN domain is in green, the RdRp in magenta, nsp7 in violet, nsp8 in light blue and nsp9 in gold. **b, c.** Coulomb density maps of the N-terminus of nsp9 from this study (**b)** and by Yan et al. (**c**) ^21^ (PDB ID:7CYQ). The NiRAN domain is shown in green and nsp9 is in gold. The arrow in (**c**) indicates additional density that likely corresponds to unmodeled Gly-Ser residues at the non-native N-terminus of nsp9. **d.** Electrostatic surface view of the NiRAN active site from 7CYQ bound to nsp9 (gold). The N-terminus and the C-terminal helix of nsp9 are shown. Electrostatic surface of nsp12 is contoured at 5 kT. **e.** Cartoon representation depicting the interactions between the nsp9 N-terminus (gold) with the β8-β9-β10 sheet in the NiRAN domain (green). Asn2 in nsp9 forms electrostatic interactions with Asp36 in the NiRAN domain and Arg733 in the RdRp domain (magenta). PDB ID 7CYQ was used. **f.** Cartoon representation depicting the interactions between the nsp9 C-terminal helix (gold) and the β8-β9-β10 sheet in the NiRAN domain (green). Interactions between Asn95/96 in nsp9 and D291 in the NiRAN domain are indicated. PDB ID 7CYQ was used. **g.** Incorporation of 5’-pRNA^LS10^ into nsp9 and deRNAylation of nsp9-pRNA^LS10^ by WT nsp12, the NiRAN mutant (D218A), the polymerase mutant (D760A), or the isolated NiRAN domain (residues 1-326; ΔRdRP). Reaction products were analysed as in **Fig. 2b. h.** Incorporation of 5’-pRNA^LS10^ into nsp9 (or the indicated mutants) by nsp12. Reaction products were analysed as in **Fig. 2b. i.** Cartoon representation of the NiRAN active site. Catalytic residues and GDP are shown as sticks, Mg^2+^ is a green sphere, and interactions are denoted by dashed lines. **j.** Relative viral yields from WT or mutant SARS-CoV-2 viruses bearing indicated mutations in nsp9 and nsp12. Data represent averages of two biological replicates. Error bars, SD. Results shown in **g** and **h** are representative of at least 2 independent experiments.

The first four residues of nsp9 extend into the NiRAN active site, forming electrostatic and hydrophobic contacts in and around a groove near the kinase-like active site (**Fig. 5d**). Asn1 of nsp9 is positioned inside of the active site, primed for transfer of 5’-pppRNA onto its N-terminus. Although the terminal NH_2_ group of nsp9 is the substrate for RNAylation, the local quality of the structures is not high enough to distinguish its exact position. We have modelled the nsp9 acceptor NH_2_ pointing towards what appears to be the phosphates of the nucleotide analogue UMP-NPP in the active site (**Fig 5b**). In the structure by Yan et al.^21^, Asn1 was assigned an opposite conformation and there are unmodeled residues (non-native N-terminus; NH_2_-Gly-Ser-) visible in the density maps, distorting local structural features (**Fig. 5c**, arrow)^27^.

Asn2 of nsp9 is in a negatively charged cleft around the NiRAN active site, and contacts Arg733, which extends from the polymerase domain and is partially responsible for positioning nsp9 (**Fig. 5e**). Both Leu4 and the C-terminal helix of nsp9 form hydrophobic interactions with a β-sheet (β8-β9-β10) in the N-lobe of the NiRAN domain (**Fig. 5e, f**). The N-terminal cap of the nsp9 C-terminal helix also forms electrostatic interactions with a negatively charged pocket on the surface of the NiRAN domain (**Fig. 5f**). Nsp12 lacking the RdRp domain (ΔRdRp; 1-326) neither RNAylates nsp9 nor processes nsp9-pRNA^LS10^ to form GpppA-RNA (**Fig. 5g**). Likewise, deleting the C-terminal helix on nsp9 (ΔC; 1-92) and Ala substitutions of Asn1 and Asn2 abolished RNAylation (**Fig. 5h**).

The NiRAN domain resembles SelO, with an RMSD of 5.7 Å over 224 Cα atoms (PDB ID: 6EAC^13^, **Extended Data Fig. 10a**). Lys73 (PKA nomenclature; K72) forms a salt bridge with Glu83 (PKA; E91) from the aC (a2) helix and contacts the phosphates (GDP in 7CYQ, or UMP-NPP in our structure; **Fig. 5i**). As expected, the “DFG” Asp218 (PKA; D184) binds a divalent cation. Interestingly, the NiRAN domain lacks the catalytic Asp (**Extended Data Fig. 1**), (PKA; D166); however, like in SelO, Asp208 is next to the metal binding Asn209 (PKA; N171) and may act as a catalytic base to activate the NH_2_ group on the N-terminus of nsp9 (**Fig. 5i**).

In canonical kinases, the β1-β2 G-loop stabilizes the phosphates of ATP^28^. In contrast, the NiRAN domain contains a β-hairpin insert (β2-β3) where the β1-β2 G-loop should be (**Extended Data Fig. 10b**). This insertion not only makes contacts with the N-terminus of nsp9, but also contains a conserved Lys (K50) that extends into the active site and stabilizes the phosphates of the bound nucleotide. Likewise, Arg116 also contacts the phosphates of the nucleotide. SelO contains a similar set of basic residues pointing into the active site that accommodate the flipped orientation of the nucleotide to facilitate AMPylation (**Extended Data Fig. 10b**). Notably, Lys73, Arg116 and Asp218 in SARS-CoV-1 nsp12 are required for viral replication^5^.

### The kinase-like residues of the NiRAN domain and the N-terminus of nsp9 are essential for SARS-CoV-2 replication

To determine the importance of the NiRAN domain and the N-terminus of nsp9 in viral replication, we used a DNA-based reverse genetics system that can rescue infectious SARS-CoV-2 (Wuhan-Hu-1/2019 isolate) expressing a fluorescent reporter^8^ (**Extended Data Fig. 11a**). We introduced single point mutations in nsp9 (N1A, N1D and N2A) and nsp12 (K73A, D218A and D760A) and quantified the virus in supernatants of producer cells by RT-qPCR to detect the viral N gene. We observed a 400 to 4000-fold reduction in viral load for all the mutants compared to WT (**Fig. 5j, Extended Data Fig. 11a).** To account for the possibility of a proteolytic defect in the mutant viral polyprotein, we tested whether the main viral protease nsp5 (M^Pro^) can cleave a nsp8-nsp9 fusion protein containing the Asn1/Asn2 mutations in nsp9. The N1D mutant failed to be cleaved by nsp5, suggesting that the replication defect observed for this mutant is a result of inefficient processing of the viral polyprotein. However, the N1A and N2A mutants were efficiently cleaved by nsp5 (**Extended Data Fig. 11b, c**). Collectively, these data provide genetic evidence that the residues involved in capping of the SARS-CoV-2 genome are essential for viral replication.

## Discussion

We propose the following mechanism of RNA capping by CoV: during transcription, the nascent 5’-pppRNA binds to the NiRAN active site, in either a *cis* (**Fig. 6a**) or *trans* **(Fig. 6b)** manner and 5’-pRNA is subsequently transferred to the N-terminus of nsp9 forming a phosphoramidate bond **(Fig. 6c, panels 1 and 2)**. The nsp13 protein produces GDP from GTP, which binds the NiRAN active site and attacks RNAylated nsp9, releasing capped RNA and regenerating unmodified nsp9 (**Fig. 6c, panels 3 and 4)**. Subsequently, nsp14 and nsp16 perform sequential N7 and 2’-O methylations, forming a fully functional ^7Me^GpppA_2’-O-Me_-RNA cap.

**Figure 6.**
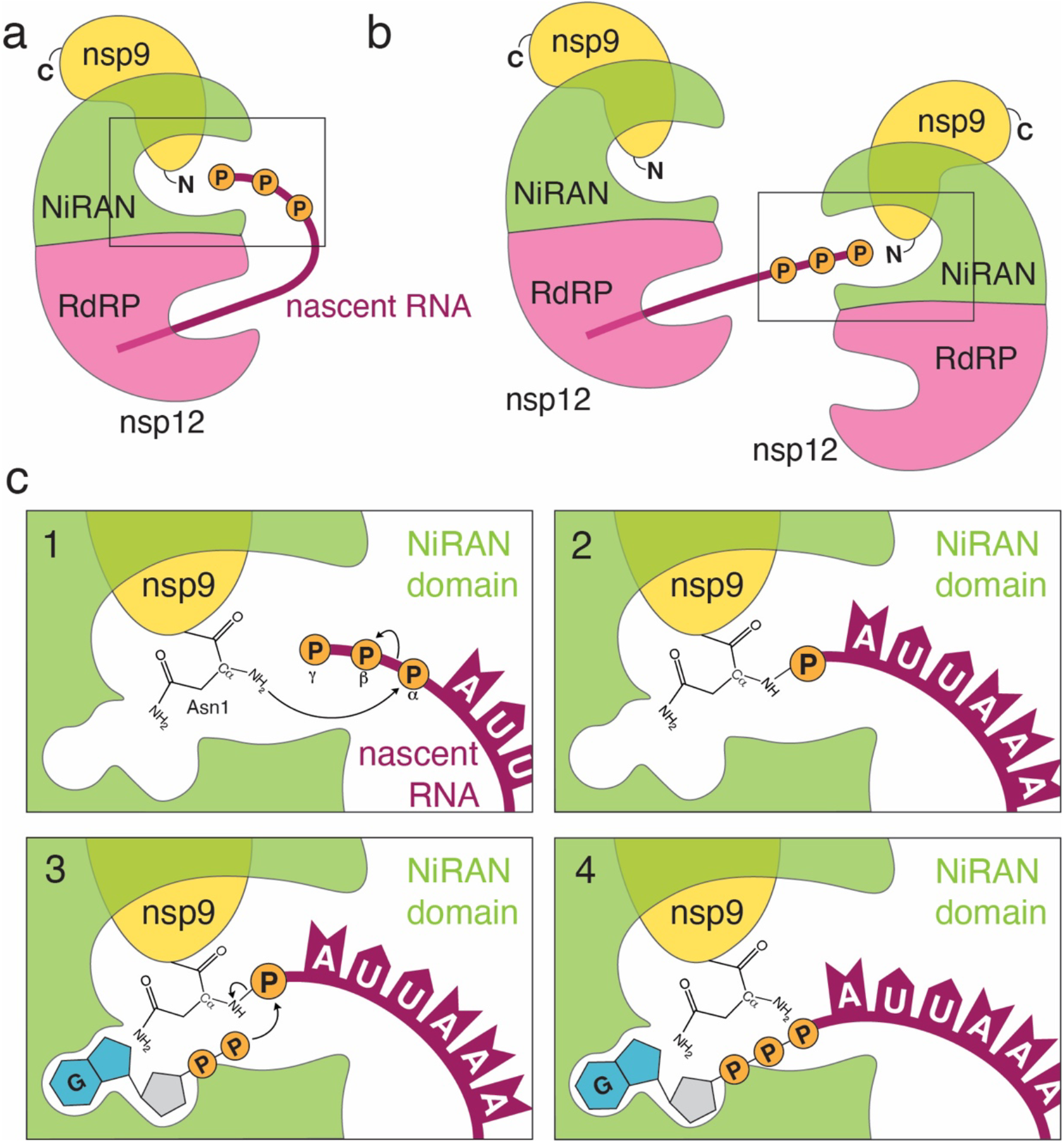
Proposed model of the SARS-CoV-2 RNA capping mechanism. **a, b.** During transcription, the nascent 5’-pppRNA binds to the NiRAN active site in either a *cis* (**a**) or a *trans* (**b**) manner. **c.** Upon binding, the N-terminus of nsp9 attacks the α-phosphate of the nascent 5’-pppRNA (**1**), forming the covalent nsp9-pRNA species and releasing PPi (**2**). Upon GDP binding to the NiRAN active site, the β-phosphate of the nsp13-generated GDP attacks the 5’-phosphate on the nsp9-pRNA (**3**), releasing capped RNA and regenerating unmodified nsp9 (**4**). Subsequent methylation events are carried out by nsp14 and nsp16 to generate the ^7Me^GpppA_2’-O-Me_-RNA cap.

SARS-CoV-2 nsp12 is thought to initiate transcription/replication starting with an NTP, or a short 5’-pppRNA primer^29^. Cryo-EM structures of the RTC suggest that the dsRNA product makes its way out of the RdRp active site in a straight line, supported by the nsp8 helical stalks^21,30,31^. In a *cis* capping model, the helical duplex with nascent 5’-pppRNA would then need to unwind, flex 90°, and extend into the NiRAN active site ~70 Å away (**Fig. 6a**). More likely, a separate RTC complex could perform capping in *trans* (**Fig. 6b**). Notably, Perry et al.^32^ propose that the nascent RNA strand is separated from the template upon passing through the proof-reading ExoN domain of nsp14 on a neighbouring RTC and threaded towards the NiRAN domain.

SARS-CoV-1 nsp13 has RNA helicase, nucleotide triphosphatase (NTPase), and RNA 5’-triphosphatase (RTPase) activities^20^. The RTPase activity implicated nsp13 in the first step of the capping mechanism; however, while nsp13 can act on 5’-pppRNA, this reaction is inhibited in the presence of cellular concentrations of ATP^20^. Thus, we favour the idea that the physiological functions for nsp13 are: **1)** to utilize the energy from ATP hydrolysis to unwind double-stranded RNA (helicase), and **2)** to hydrolyse GTP to GDP, which can then act as an acceptor for 5’-pRNA in the NiRAN-catalysed capping reaction.

The SARS-CoV-2 capping mechanism is reminiscent of the capping mechanism used by VSV, although there are some differences. The VSV large (L) protein is a multifunctional enzyme that carries out RdRp, PRNTase, and methyltransferase activities to form the cap^24,33,34^. During the reaction, 5’-pRNA is transferred to a conserved His within the PRNTase domain, which adopts a unique α-helical fold that is distinct from that of protein kinases^25^. The presence of two different enzymatic mechanisms of capping, proceeding via covalent protein-RNA intermediates, in *Mononegavirales* and in *Nidovirales* is an example of convergent evolution.

Consistent with other reports^16,27^, we observed NiRAN-catalysed NMPylation of nsp9 (**Fig. 1b, Extended Data Fig. 3**). While our results do not necessarily preclude a biologically relevant function for nsp9 NMPylation, it is worth noting that this modification is reversible in the presence of PPi^27^ (**Fig. 3e**). PPi is produced during the RdRp reaction, making the stability of NMPylated nsp9 difficult to envision in vivo. By contrast, RNAylated nsp9 was not reversible in the presence of PPi. Thus, RNAylation is likely the physiologically relevant modification of nsp9 during viral RNA capping.

Recent work suggested that the NiRAN domain is a GTase that transfers GMP from GTP to 5’-ppRNA, forming a GpppA-RNA cap intermediate^3,21^. In our efforts to reproduce these results, we failed to detect nsp12-dependent GpppA cap formation by TLC (**Extended Data Fig. 12a**) or by Urea-PAGE analysis of the RNA (**Extended Data Fig. 12b**), in contrast to our control, in which the Vaccinia capping enzyme efficiently generated GpppA-RNA. Because nsp13 and the Vaccinia capping enzyme can hydrolyse GTP to GDP^20^, the cap reported previously^21^ appears to be GDP formed from nsp13- and Vaccinia capping enzyme-dependent hydrolysis of GTP.

In summary, we have defined the mechanism by which SARS-CoV-2 caps its genome and have reconstituted this reaction in vitro using non-structural proteins encoded by SARS-CoV-2. Our results uncover new targets for the development of antivirals to treat COVID-19 and highlight the catalytic adaptability of the kinase domain.

