## Supplemental Materials for "The mechanism of RNA capping by SARS-CoV-2"

### Extended Data Figures

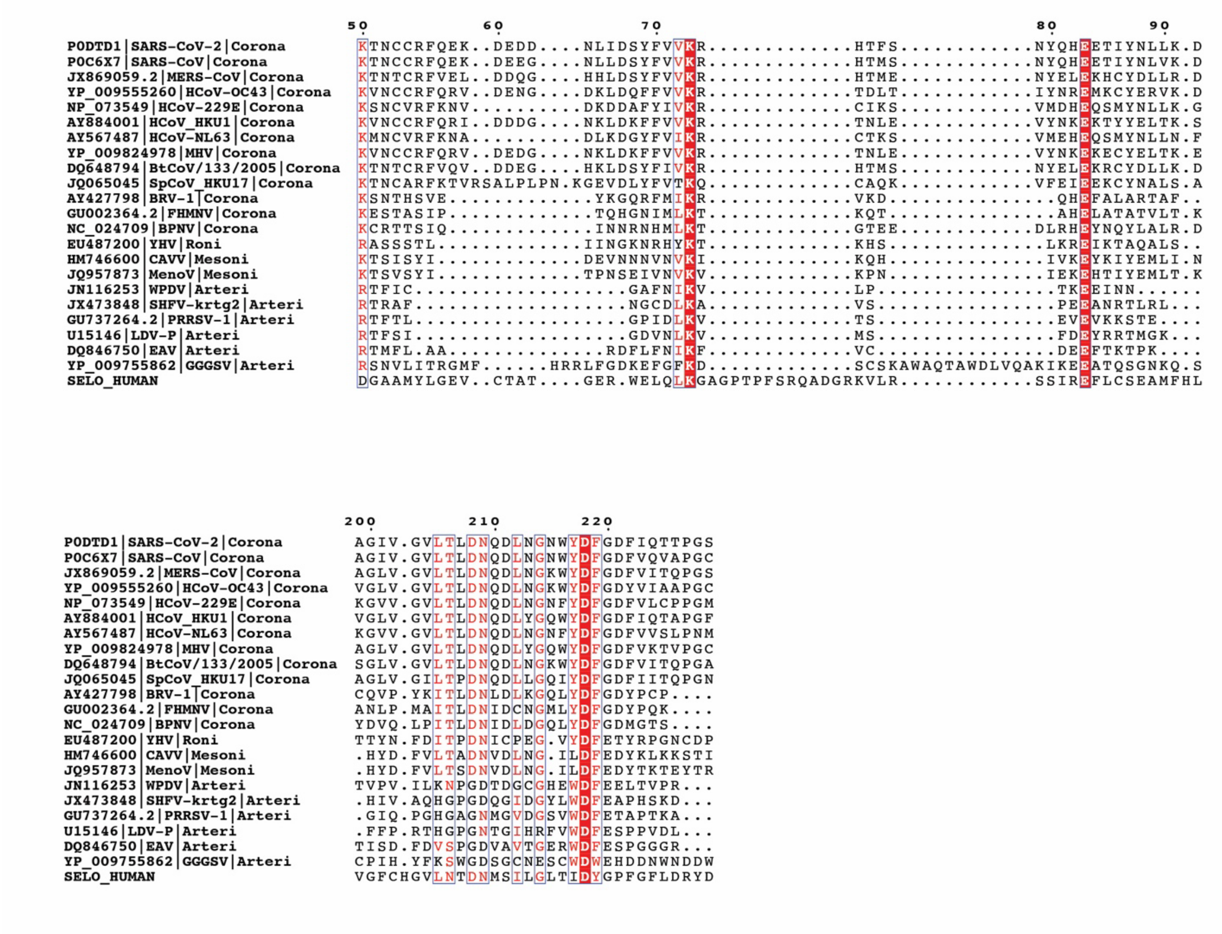


**Extended Data Fig. 1. Sequence alignment of the NiRAN domain reveals similarity to the pseudokinase selenoprotein-O (SelO).** Multiple sequence alignment highlighting conserved kinase-like active site residues in the NiRAN domain among several CoVs, other selected *Nidovirales* (Arteri-, Mesoni- and Roniviruses) and the human SelO pseudokinase. Top: amino acid sequence surrounding the Lys-Glu ion pair.  Bottom: and the amino acid sequence surrounding the active site and the "DFG" motif.

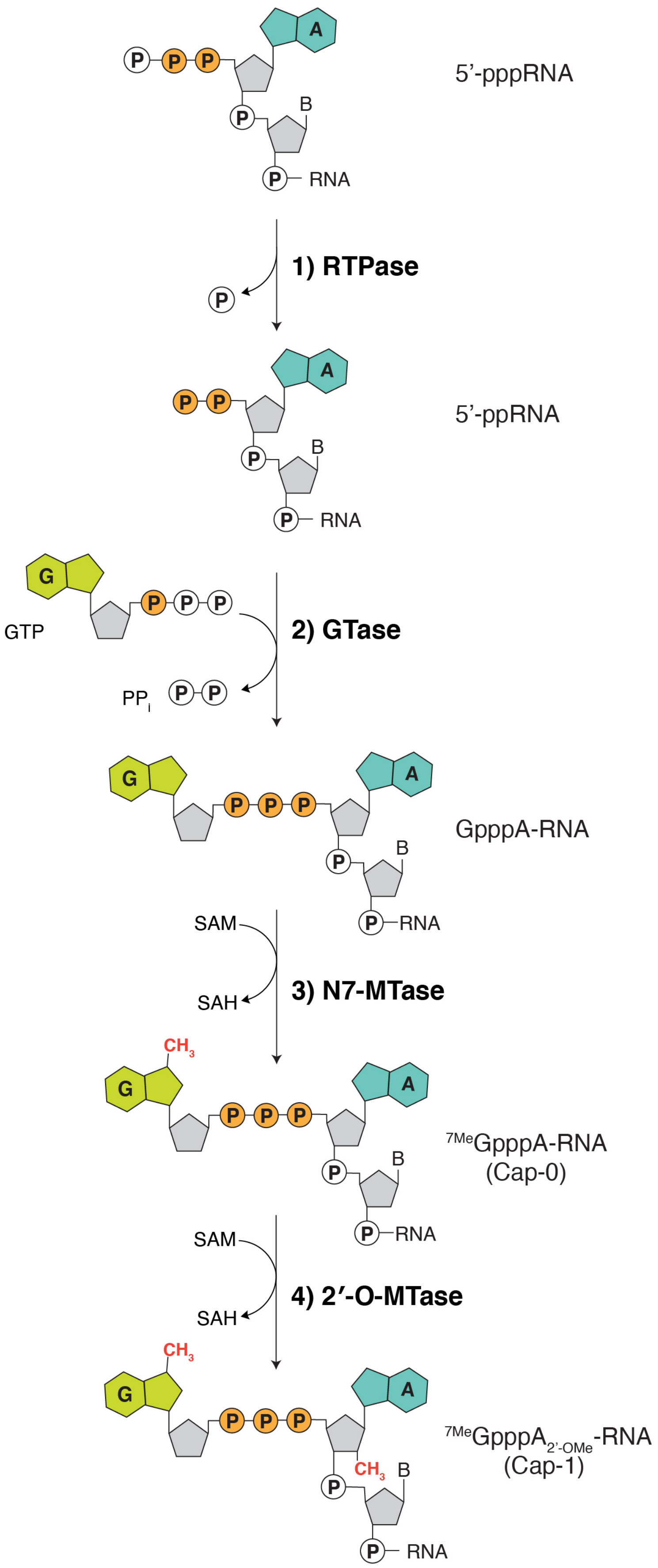

P P P
A
5’-pppRNA
B
P RNA
1) RTPase
P
P P
A
5’-ppRNA
B
P RNA
G
GTP
P P P
2) GTase
PPi
P P
G
P P P
A
GpppA-RNA
B
P RNA
SAM
SAH
3) N7-MTase
CH3
G
P P P
A
7MeGpppA-RNA
(Cap-0)
B
P RNA
SAM
SAH
4) 2′-O-MTase
CH3
G
P P P
A
7MeGpppA2’-OMe-RNA
(Cap-1)
CH3
B
P RNA


9

**Extended Data Fig. 2. The canonical eukaryotic mRNA capping mechanism.** The $^{7Me}GpppA_{2'OMe}$ cap on eukaryotic RNA is formed co-transcriptionally by four enzymes: **1)** an RNA triphosphatase (RTPase), which removes the γ-phosphate from the nascent 5'-triphosphorylated RNA (5'-pppRNA) to yield a 5'-diphosphorylated RNA (5'-ppRNA); **2)** a guanylyltransferase (GTase), which transfers the GMP moiety from GTP to the 5'-ppRNA to form the core cap structure GpppN-RNA; **3)** a (guanine-N7)-methyltransferase (N7-MTase), which methylates the cap guanine at the N7 position; and **4)** a (nucleoside-2′-*O*)-methyltransferase (2′-O-MTase), which methylates the ribose-2'-OH position on the first nucleotide of the RNA. B denotes any base; GTP, Guanosine triphosphate; GDP, Guanosine diphosphate; $PP_i$, pyrophosphate; SAM, S-Adenosyl methionine.

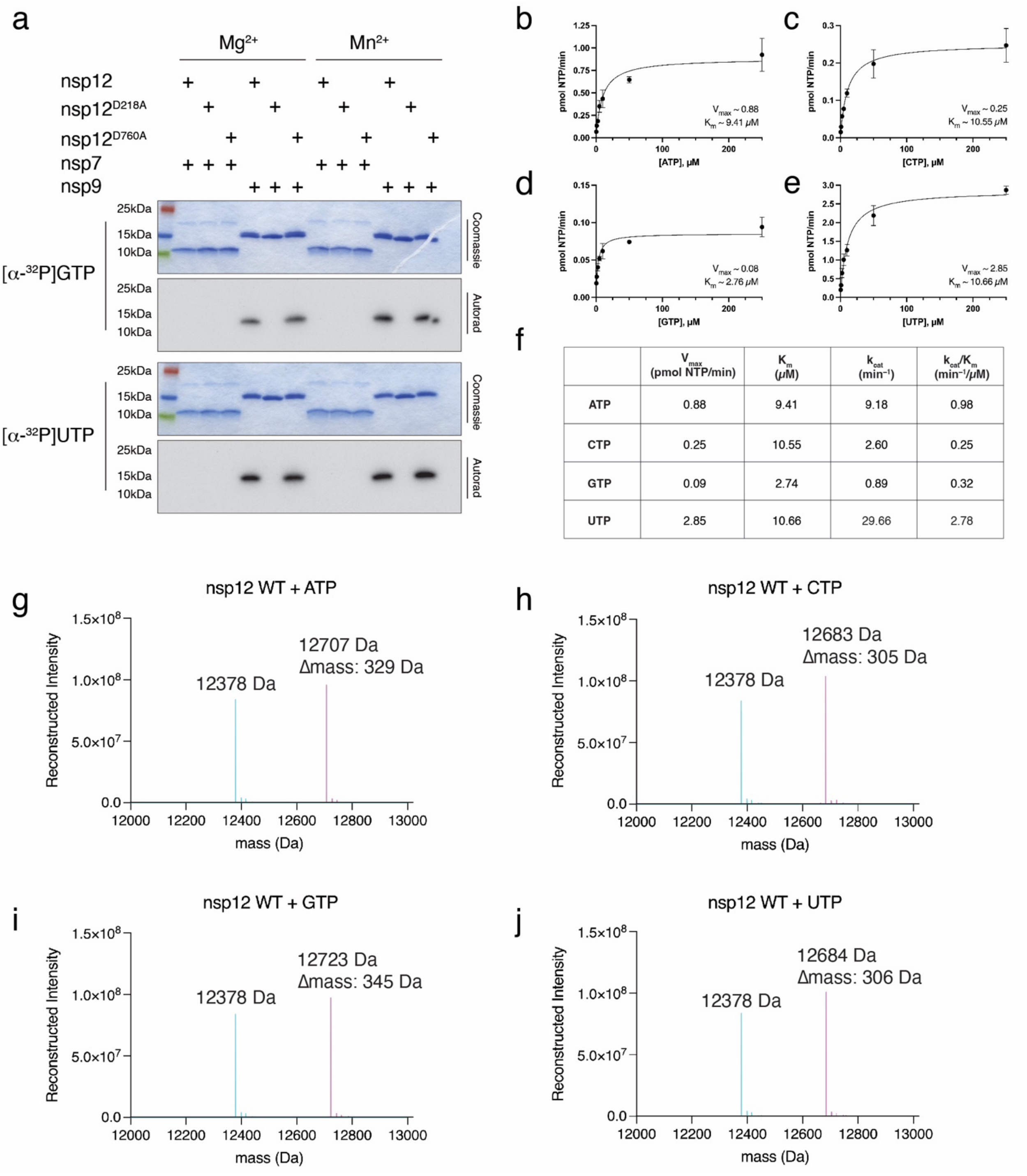


| | $V_{max}$ (pmol NTP/min) | $K_m$ (μM) | $k_{cat}$ (min$^{-1}$) | $k_{cat}/K_m$ (min$^{-1}$/μM) |
|---|---|---|---|---|
| ATP | 0.88 | 9.41 | 9.18 | 0.98 |
| CTP | 0.25 | 10.55 | 2.60 | 0.25 |
| GTP | 0.09 | 2.74 | 0.89 | 0.32 |
| UTP | 2.85 | 10.66 | 29.66 | 2.78 |

**Extended Data Fig. 3. The NiRAN domain NMPylates nsp9. a.** Incorporation of α-$^{32}$P from [α-$^{32}$P]GTP or [α-$^{32}$P]UTP into nsp7 or nsp9 by WT nsp12, the NiRAN mutant (K73A, D218A), or the polymerase mutant (D760A). Reactions were performed in the presence of $Mg^{2+}$ or $Mn^{2+}$ and the products were resolved by SDS-PAGE and visualized by Coomassie staining (top) and autoradiography (bottom). **b-e**. Kinetic analysis depicting the concentration dependence of (**b**) ATP, (**c**) CTP, (**d**) GTP, or (**e**) UTP on the rate of nsp9 NMPylation by the NiRAN domain. $K_m$ and $V_{max}$ are indicated on the insets. Plots shown are the mean and SD of triplicate reactions. **f.** Summary of $K_m$, $V_{max}$, $k_{cat}$, and $k_{cat}/K_m$ values for each NTP. **g-j**. Intact mass LC/MS spectra of unmodified nsp9 (*cyan)* overlayed with NMPylated nsp9 (*pink*) following incubation with WT nsp12 and (**g**) ATP, (**h**) CTP, (**i**) GTP, or (**j**) UTP. The observed masses are shown in the insets. The theoretical mass of unmodified nsp9 is 12378.2 Da and the theoretical increase in mass with the addition of each NMP is as follows: AMP, 329 Da; CMP, 305 Da; GMP, 345 Da; UMP, 306 Da.

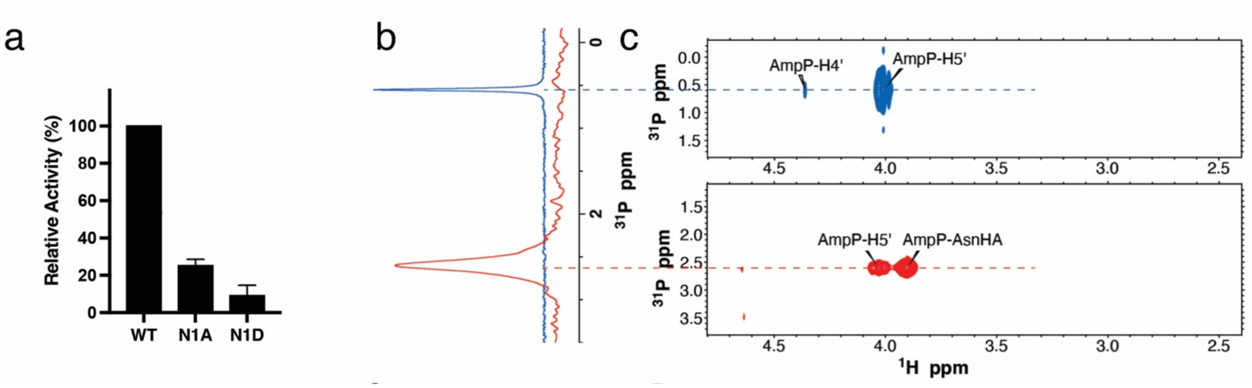


**Extended Data Fig. 4. The NiRAN domain NMPylates nsp9 on the N-terminus. a.** Quantification of reaction products from **Fig. 1c** depicting the relative NiRAN-dependent UMPylation activity towards nsp9 or the indicated mutants. Radioactive gel bands were excised and quantified by scintillation counting. **b.** 1D 31P spectrum of AMP-nsp9 (*red*) and AMP (*blue*) recorded in the same buffer as reference. **c.** 2D 1H, 31P-HSQC spectra of AMP (*top, blue*) and AMP-nsp9 (*bottom, red*).

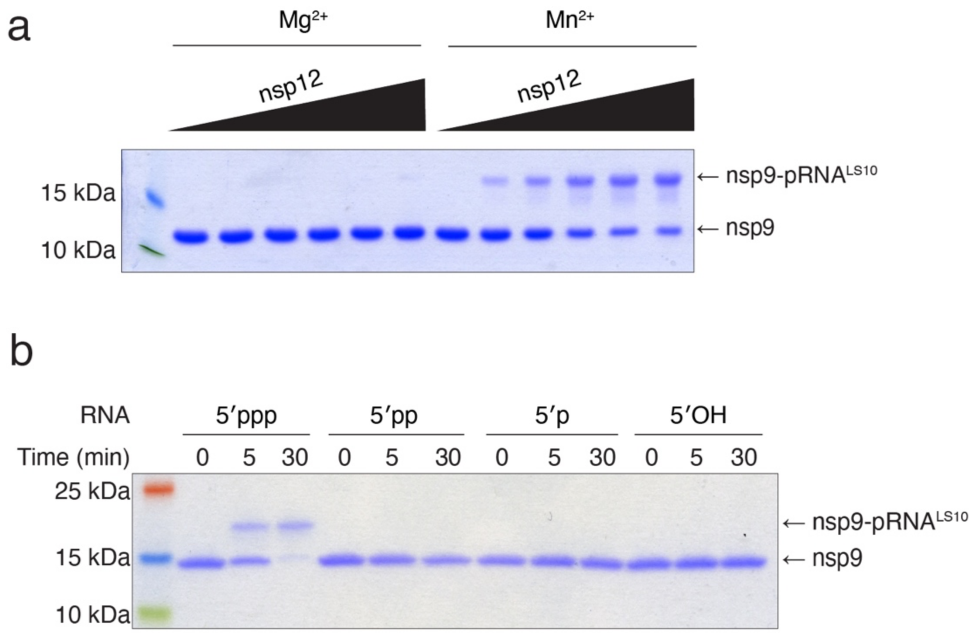


**Extended Data Fig. 5. Characterization of NiRAN RNAylation activity. a.** Incorporation of RNA into nsp9 by nsp12 (0-4 μM) in the presence of $Mg^{2+}$ or $Mn^{2+}$. Reaction products were analysed as in **Fig. 2b**. **b.** Incorporation of RNA with the indicated 5' ends into nsp9 by nsp12. Reaction products were analysed as in **Fig. 2b**.

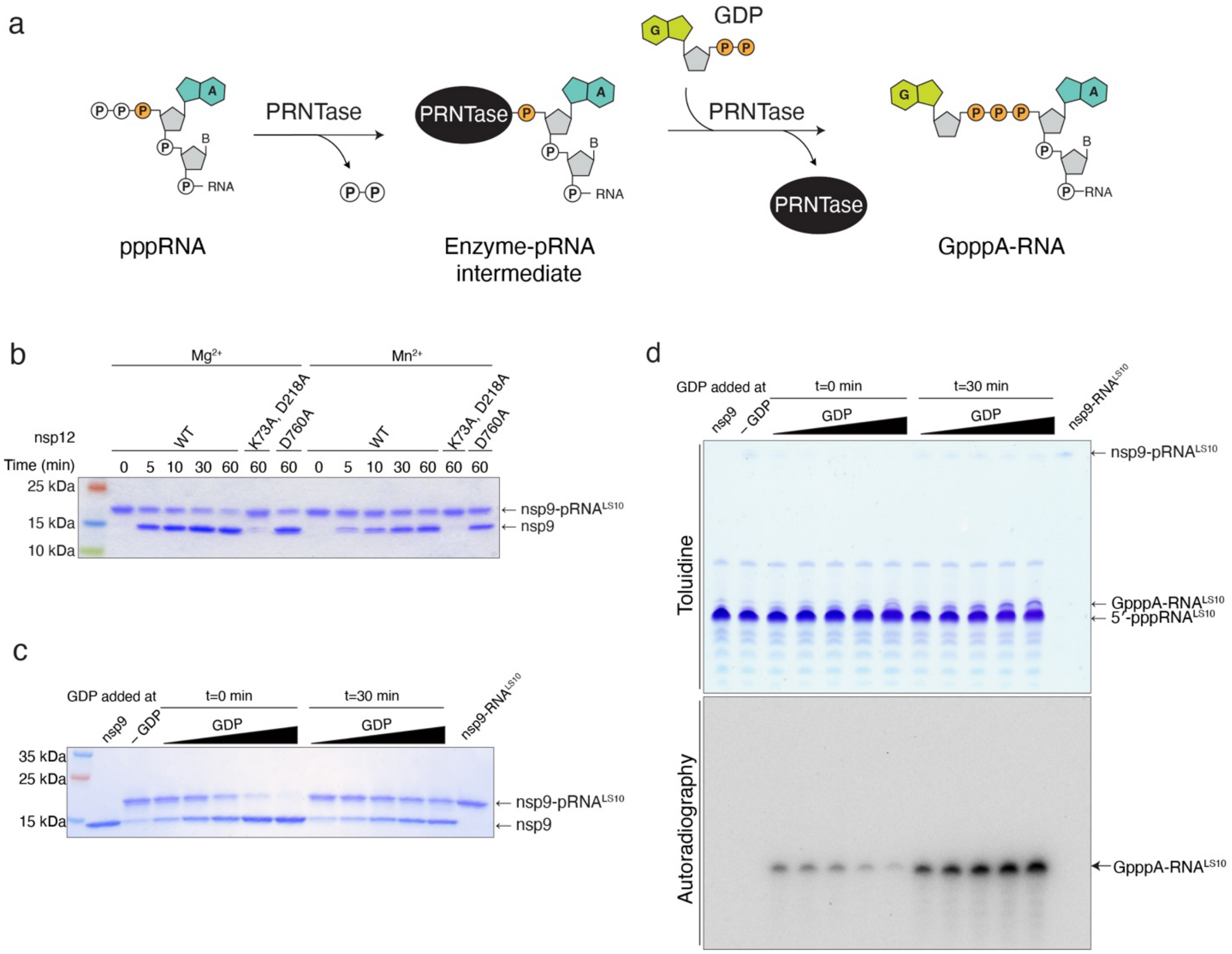


**Extended Data Fig. 6. Characterization of nsp12 NiRAN GDP-PRNTase activity. a.** Schematic representation depicting the mechanism of GpppA-RNA formation by vesicular stomatitis virus (VSV) polyribonucleotidyltransferase (PRNTase) enzyme. **b.** Time-dependent deRNAylation of nsp9-pRNA$^{LS10}$ by WT nsp12, the NiRAN mutant (K73A, D218A), or the polymerase mutant (D760A) in the presence of GDP and either $Mg^{2+}$ or $Mn^{2+}$. Reaction products were analysed as in **Fig. 2b**. **c, d.** NiRAN-catalysed capping reactions depicting the inhibitory effect of GDP on RNAylation. Nsp9 was incubated with excess 5'-pppRNA$^{LS10}$ in presence of nsp12 with no GDP (-GDP), or increasing concentrations (6.25-100 µM) of [$^{32}$P]GDP added either at time zero (t =0 min), or after the RNAylation reaction was allowed to proceed for 30 minutes (t=30 min). Reaction products were analysed by SDS-PAGE **(c)**, and Urea-PAGE **(d)**.

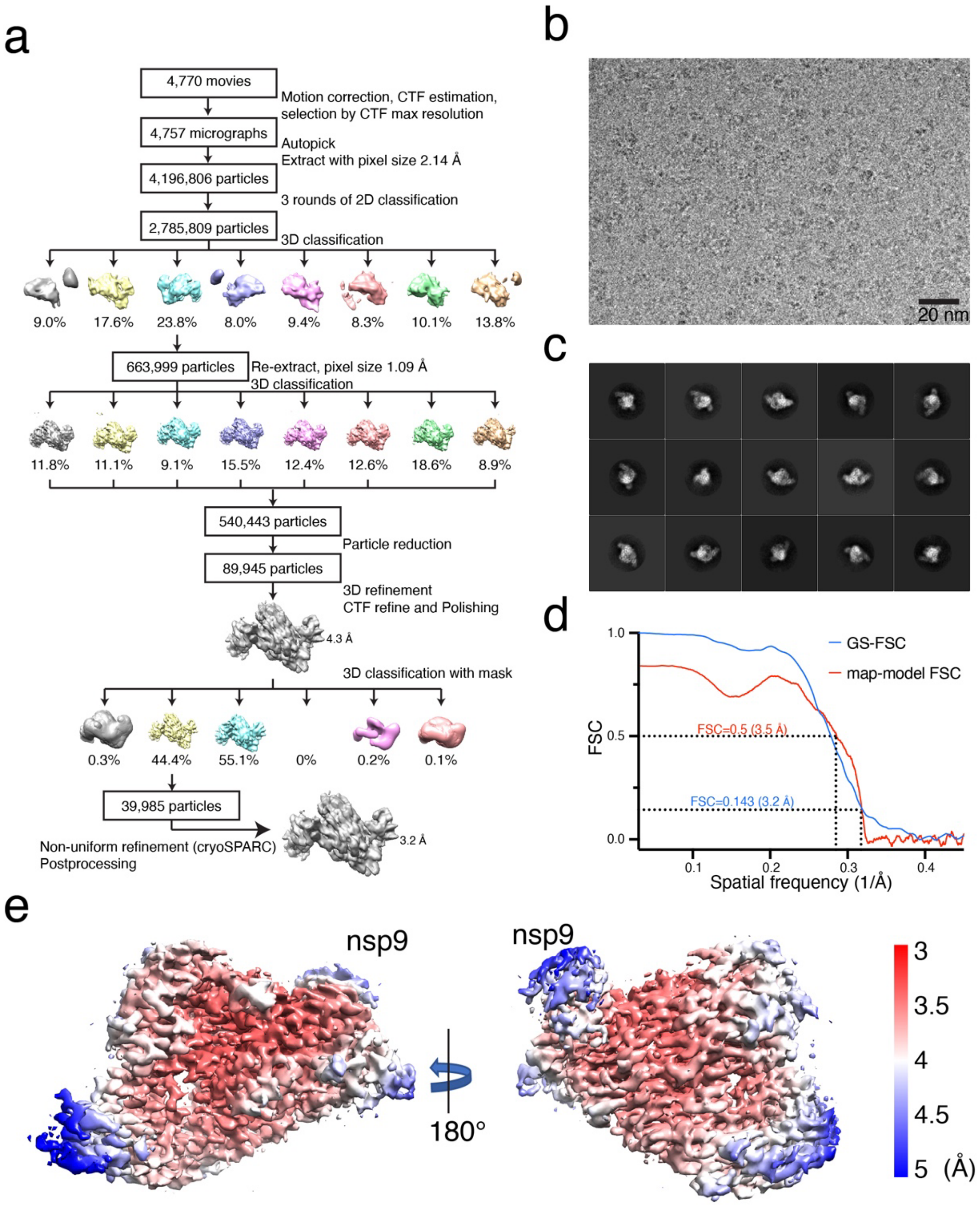


**Extended Data Fig 7. Cryo-EM analysis of the nsp7/8/9/12 complex. a.** Flow chart representing data processing for the nsp7/8/9/12 complex. **b.** A representative micrograph of the nsp7/8/9/12 complex grids. **c.** Representative 2D classes generated by RELION 2D-classification. **d.** Gold-standard FSC curve (blue), and map-model FSC curve (red). Curves were generated by cryoSPARC and Phenix suite, respectively. **e.** Local resolution of the nsp7/8/9/12 complex calculated by RELION from final cryoSPARC half-maps. Position of nsp9 is indicated.

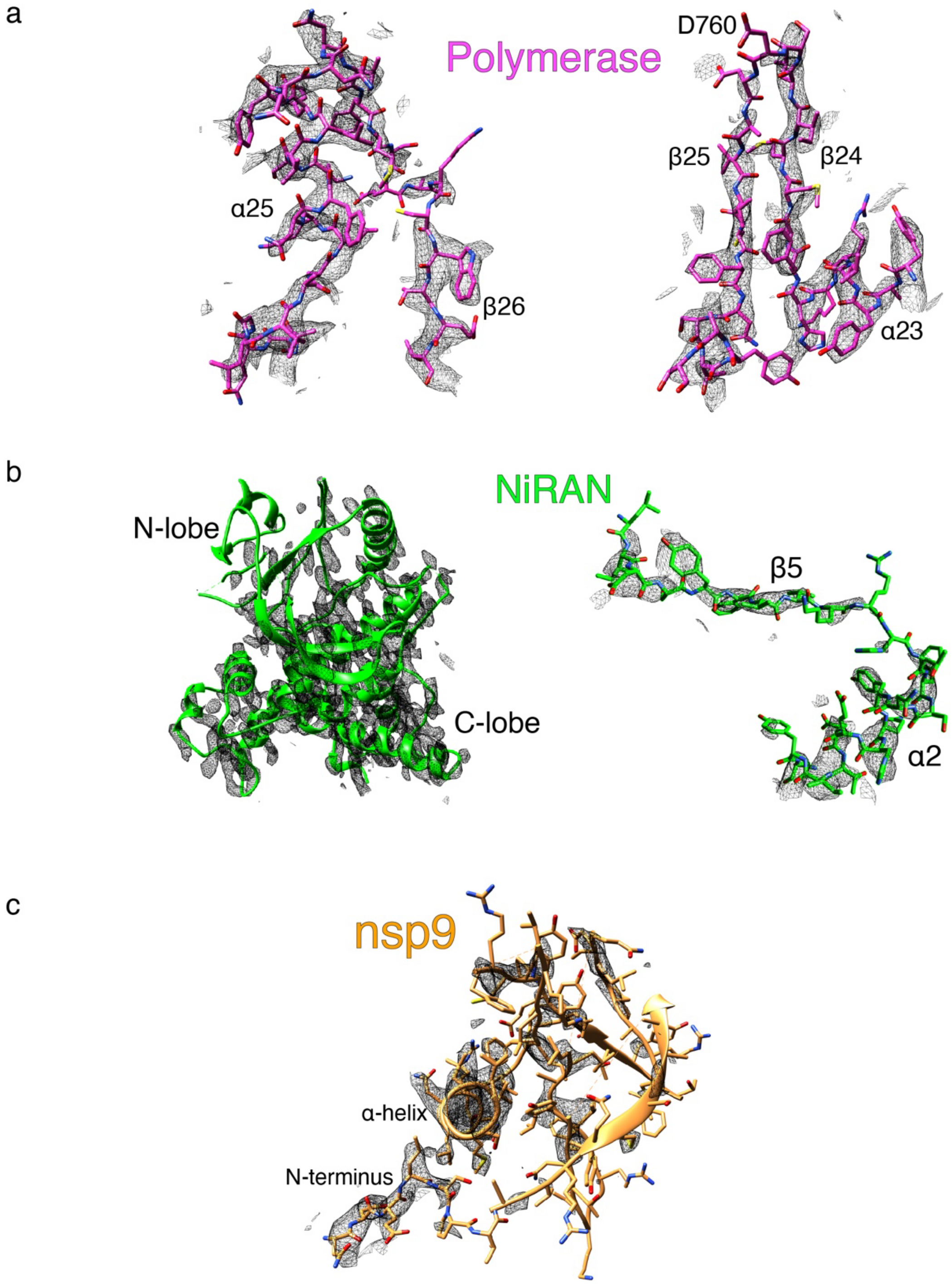


**Extended Data Fig 8. Exemplary cryo-EM density (black mesh)** on (**a**) Polymerase (Magenta), (**b**) NiRAN (green) and (**c**) nsp9 (gold). Note weaker density in the N-lobe of the kinase-like NiRAN domain (left panel of **b**, compare top to bottom of the image), and poor density in nsp9 (**c**), in areas not in direct contact with the NiRAN domain.

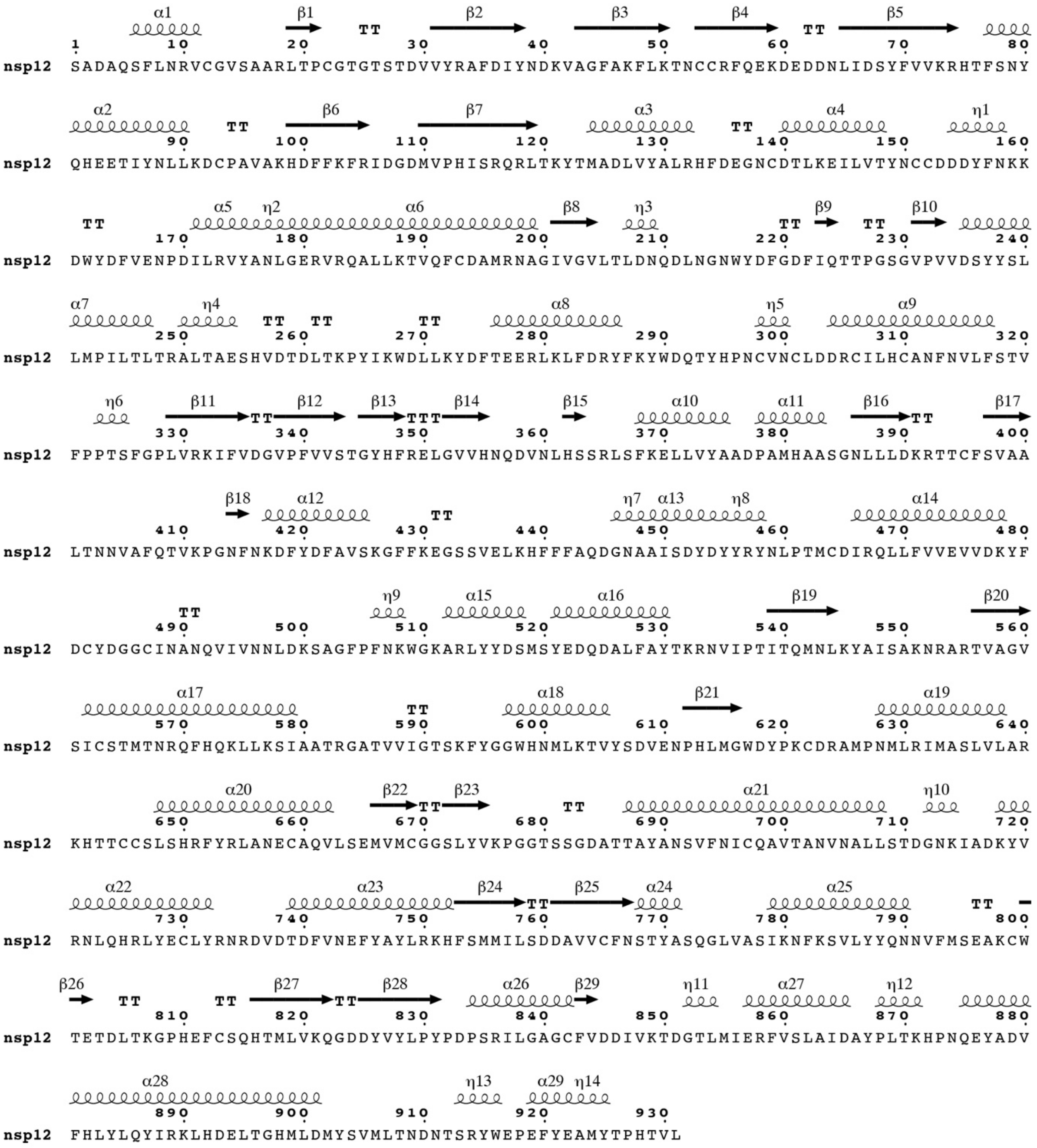


**Extended Data Fig. 9. Secondary structure of nsp12.** The secondary structural elements in nsp12 (from PDB ID: 7CYQ) are shown.

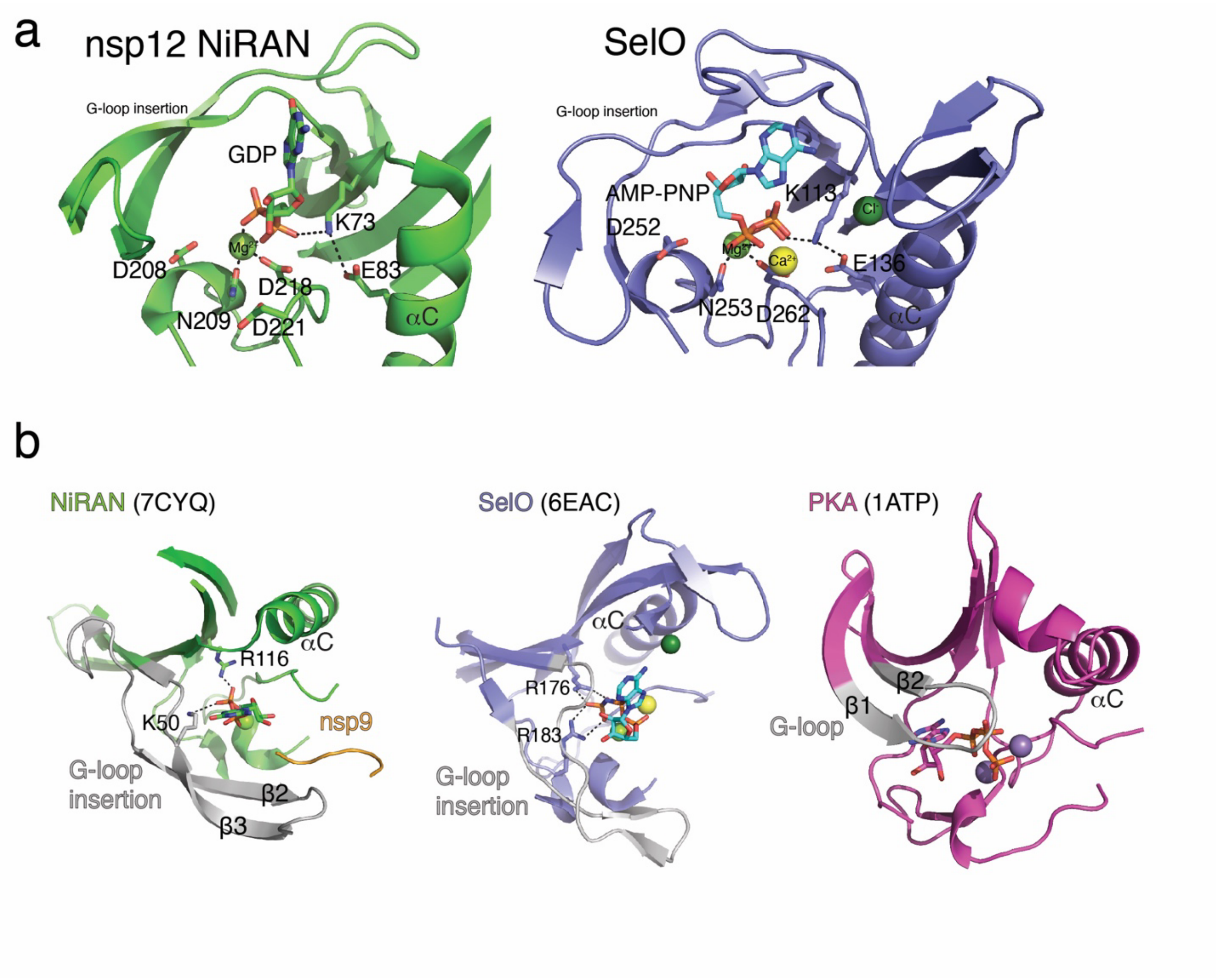


**Extended Data Fig. 10. Comparison of the kinase-like domains of nsp12 and SelO. a.** Cartoon representation comparing the NiRAN active site catalytic residues (green) to the active site residues in SelO (purple). The divalent cations are shown as spheres. **b.** Comparison of the Gly-rich loop regions in NiRAN (PDB ID: 7CYQ, left), SelO (PDB ID: 6EAC), and the canonical kinase PKA (PDB ID: 1ATP, right). Green sphere – $Mg^{2+}$, dark green sphere – Chloride, yellow sphere – calcium, violet sphere – $Mn^{2+}$.

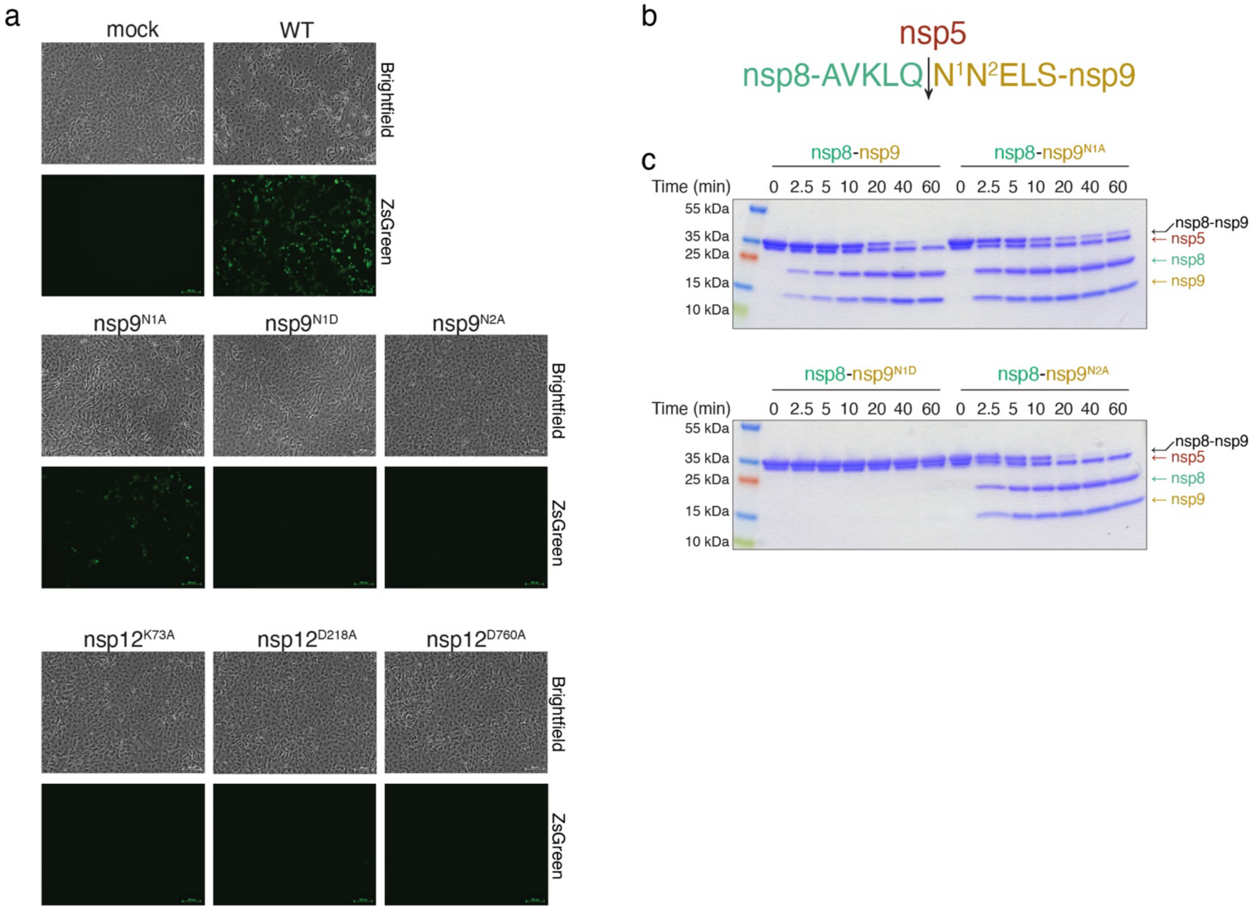


**Extended Data Fig. 11. Genetic insights into RNA capping by the NiRAN domain. a.** Microscopy images showing brightfield (upper) or fluorescence-based images (ZsGreen; lower) of SARS-CoV-2-ZsGreen production in VeroE6-C1008-TMPRSS2 cells. Mock-transfected panels were incubated with transfection reagents lacking DNA. The mutations engineered into either nsp9 or nsp12 are indicated above each set of images. Data represent one set of images from two independent biological replicates. Scale bars, 100 µm. **b.** The amino acid sequence between nsp8 (green) and nsp9 (gold) depicting the cleavage site for the nsp5 (dark red) protease. N1 and N2 of nsp9 are highlighted. The arrow denotes the location of cleavage. **c.** Time-dependent proteolysis of the nsp8-nsp9 fusion protein by nsp5. Reaction products were separated by SDS PAGE and visualized by Coomassie staining.

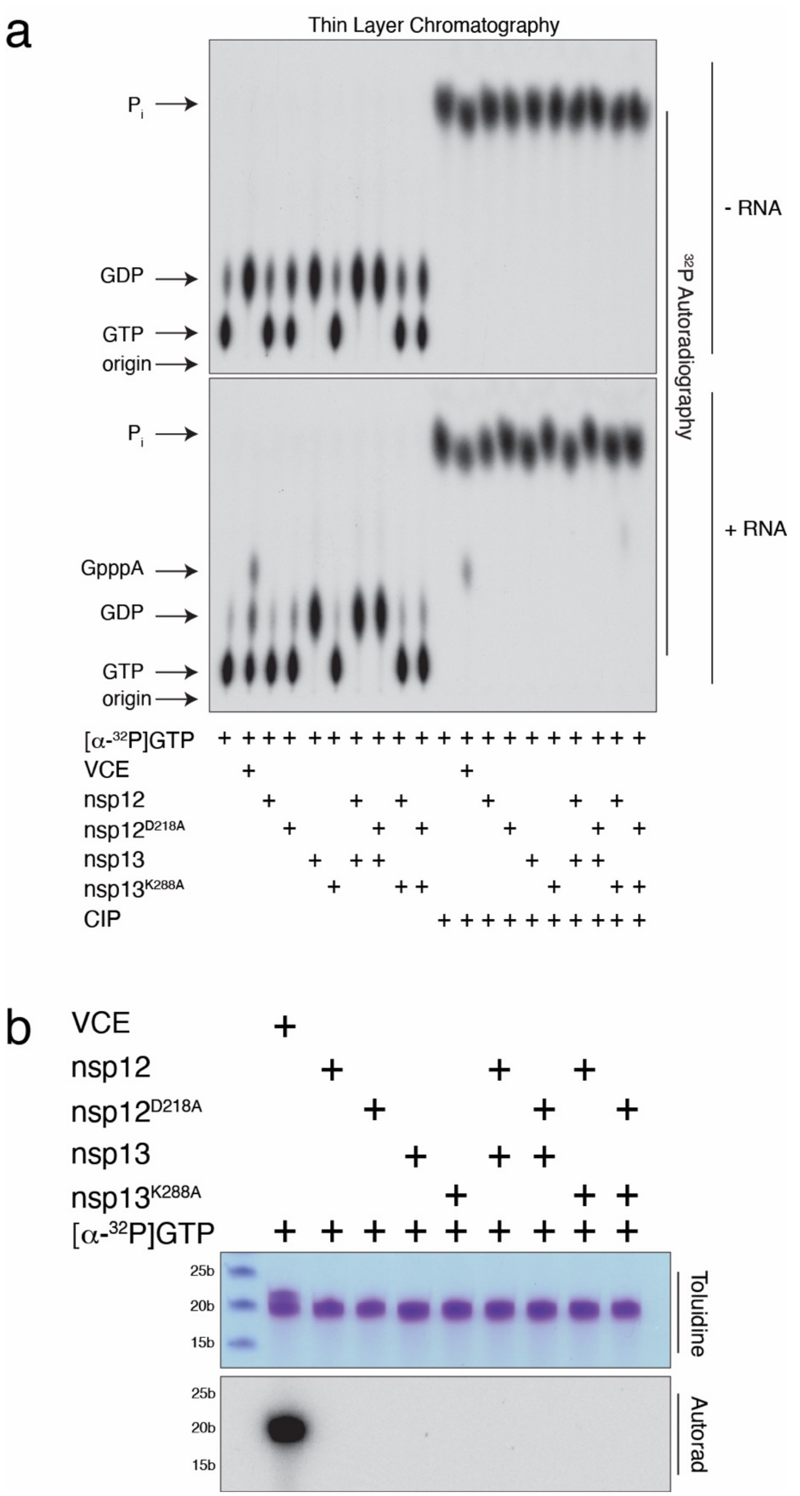


**Extended Data Fig. 12. The NiRAN domain does not act as a GTase**. **a.** Thin-layer chromatograms depicting the reaction products resulting from the incubation of [α-$^{32}$P]GTP with nsp12 or the inactive NiRAN mutant (D218A). Reactions were performed as described in [21] with (*lower*) or without (*upper*) 5'-pppRNA$^{A19C}$ and included nsp13 or the inactive mutant (K288A) as indicated. Vaccinia capping enzyme (VCE) was used as a positive control. Reaction products were digested with nuclease P1, then treated with or without calf intestinal alkaline phosphatase (CIP) and analysed by PEI-cellulose thin-layer chromatography (TLC) followed by autoradiography. The positions of the origin and standard marker compounds are indicated. **b.** RNA products from (**a**) were analysed by TBE Urea-PAGE and visualized by toluidine blue O staining (*upper*) and autoradiography (*lower*). Markers indicate RNA size by base length.

### Extended Data Table 1. RNAs used in this study

| RNA | Sequence |
|---|---|
| 5′-pppRNA$^{\mathrm{A19C}}$ [21] | [ppp]ACCCCCCCCCCCCCCCCCCC |
| 5′-pppRNA$^{\mathrm{LS10}}$ | [ppp]AUUAAAGGUU |
| 5′-pppRNA$^{\mathrm{LS2}}$ | [ppp]AU |
| 5′-pppRNA$^{\mathrm{LS3}}$ | [ppp]AUU |
| 5′-pppRNA$^{\mathrm{LS4}}$ | [ppp]AUUA |
| 5′-pppRNA$^{\mathrm{LS5}}$ | [ppp]AUUAA |
| 5′-pppRNA$^{\mathrm{LS6}}$ | [ppp]AUUAAG |
| 5′-pppRNA$^{\mathrm{LS20}}$ | [ppp]AUUAAAGGUUUAUACCUUCC |
| 5′-pppRNA$^{\mathrm{LS10_A1C}}$ | [ppp]CUUAAAGGUU |
| 5′-pppRNA$^{\mathrm{LS10_A1G}}$ | [ppp]GUUAAAGGUU |
| 5′-pppRNA$^{\mathrm{LS10_A1U}}$ | [ppp]UUUAAAGGUU |
| 5′-pppRNA$^{\mathrm{LS10_U2G}}$ | [ppp]AGUAAAGGUU |

**Extended Data Table 2. Data collection and refinement statistics.**

| | |
|---|---|
| Magnification | 81,000 |
| Voltage (kV) | 300 |
| Electron exposure ($e^{-}$Å$^{-2}$) | 54 |
| Defocus range (μm) | -1.0 to -2.5 |
| Pixel size (Å) | 1.09 |
| Symmetry imposed | C1 |
| Initial particle images (no.) | 4,196,806 |
| Final particle images (no.) | 39,985 |
| Map resolution (Å) | 3.2 |
| FSC threshold | 0.143 |
| **Refinement** | |
| Initial model used (PDB code) | - |
| Model resolution (Å) | 3.5 |
| FSC threshold | 0.5 |
| Map sharpening *B* factor (Å$^{-2}$) | -37 |
| Nonhydrogen atoms | 9,215 |
| Protein residues | 1150 |
| Ligands | 4 |
| B factors (Å$^{-2}$) | |
| Protein | 67.4 |
| Ligands | 75.9 |
| R.m.s. deviations | |
| Bond lengths (Å) | 0.002 |
| Bond angles (˚) | 0.453 |
| MolProbity score | 1.50 |
| Clashscore | 3.12 |
| Poor rotamers (%) | 0 |
| Favored (%) | 94.1 |
| Allowed (%) | 5.9 |
| Disallowed (%) | 0 |

**Extended Data Table 3. Oligonucleotides used in the SARS-CoV-2 infection experiments**

| Oligo | Sequence | Use |
|---|---|---|
| PacI Forward | GGTTGAAGCAGTTAATTAAAGTTACACTTGTG | Fragment 1 and 3 |
| 1NtoA Reverse | GCAACAGGACTAAGCTCATTAGCCTGTAATTTGACAGC | Fragment 1 |
| 1NtoD Reverse | GCAACAGGACTAAGCTCATTGTCCTGTAATTTGACAGC | Fragment 1 |
| 2NtoA Reverse | GCAACAGGACTAAGCTCAGCATTCTGTAATTTGACAGC | Fragment 1 |
| 73KtoA Reverse | GAGAAAGTGTGTCTCGCAACTACAAAGTAAG | Fragment 1 |
| MluI Reverse | CCTAAGTTGGCGTATACGCGTAATATATCTGGG | Fragment 2 and 3 |
| 1NtoA Forward | GCTGTCAAATTACAGGCTAATGAGCTTAGTCCTGTTGC | Fragment 2 |
| 1NtoD Forward | GCTGTCAAATTACAGGACAATGAGCTTAGTCCTGTTGC | Fragment 2 |
| 2NtoA Forward | GCTGTCAAATTACAGAATGCTGAGCTTAGTCCTGTTGC | Fragment 2 |
| 73KtoA Forward | CTTACTTTGTAGTTGCGAGACACACTTTCTC | Fragment 2 |

### Acknowledgments

We thank members of the Tagliabracci laboratory for discussions, David Karlin for notifying us of the similarity between SelO and the NiRAN domain, Brenden Park for help with kinetics, Andrew Lemoff for help with intact mass analysis, and Jessica Kilgore, Noelle Williams, and the resources available in the UTSW Preclinical Pharmacology Core for detection and quantitation of GpppA. We thank Sam Wilson and Suzannah Rihn for the SARS-CoV-2 infectious clone and for technical guidance and the Structural Biology Laboratory and the Cryo-Electron Microscopy Facility at UT Southwestern Medical Center which are partially supported by grant RP170644 from the Cancer Prevention & Research Institute of Texas (CPRIT) for cryo-EM studies. A portion of this research was supported by the W. M. Keck Foundation Medical Research Grant (VST, KP, JS), the National Institutes of Health grant R01GM135189 (VST), 1DP1AI158124 (JWS), a Welch Foundation Grant I-1911 (VST), a Life Sciences Research Foundation Fellowship (GH), a Polish National Agency for Scientific Exchange scholarship PPN/BEK/2018/1/00431 (KP). J.W.S. is a Burroughs Wellcome Fund Investigator in the Pathogenesis of Infectious Disease. V.S.T. is a Howard Hughes Medical Institute Investigator, the Michael L. Rosenberg Scholar in Medical Research, a CPRIT Scholar (RR150033), and a Searle Scholar.

### Author contributions

G.J.P., A.O., G.H., J.W.S., and V.S.T. designed the experiments; G.J.P., A.O., G.H., J.L.E., A.M., and V.S.T. conducted experiments; G.J.P. discovered the RNAylation and GDP-PRNTase activity of the NiRAN domain; A.O. Z.C. and Y.L. performed the cryo-EM; A.O. performed methylation experiments; G.H. performed GTase and nsp9 NMPylation experiments; A.M. performed intact mass data analysis; M.T. and K.H.W. performed NMR experiments; K.P. performed the bioinformatics; G.J.P., A.O., and G.H. purified proteins; G.J.P., A.O. and V.S.T. performed cloning and site directed mutagenesis; J.L.E. performed infectious SARS-CoV-2 experiments; and G.J.P., A.O., K.P., J.W.S., and V.S.T. wrote the manuscript with input from all authors.

### Competing interests

The authors declare no competing interests. Data and materials availability:

### Materials & Correspondence

Correspondence and material requests should be addressed to V.S.T.

The atomic coordinates have been deposited in the Protein Data Bank with accession code 7THM.

### Methods

Chemicals and reagents

Ampicillin sodium (A9518), ATP (A2383), ADP (A2754), chloramphenicol (C0378), CTP (C1506), CDP (C9755), dithiothreitol (DTT; D0632), EDTA (E5134), GTP (G8877), GDP (G7127), imidazole (I2399), IPTG (I5502), kanamycin sulfate (K1377), 2-mercaptoethanol (BME, M3148), Brilliant blue R (B0149), magnesium chloride ($MgCl_2$; M2670), manganese (II) chloride tetrahydrate ($MnCl_2$; M3634), PEI-cellulose TLC plates (Z122882), potassium chloride (KCl; P9541), pyrophosphate (221368), Urea (U6504), UTP (U6625), UDP (94330), were obtained from MilliporeSigma (St. Louis, MO). Q5 DNA polymerase (M0492L), all restriction enzymes used for cloning, Proteinase K (P8107S), Yeast Inorganic Pyrophosphatase (M2403), Quick CIP (M0525S), Nuclease P1 (M0660S), Vaccinia Capping System (M2080S), mRNA Cap 2'-O-Methyltransferase (M0366S), G(5′)ppp(5′)A RNA Cap Structure Analog (GpppA; S1406L), and m7G(5′)ppp(5′)A RNA Cap Structure Analog (m7GpppA; S1405S) were all obtained from New England Biolabs (Ipswich, MA). Acetic acid (A38-212), RNAse inhibitor (N8080119), 2X TBE-Urea Sample Buffer (LC6876), and isopropanol (42383) were all obtained from Thermo Fisher Scientific (Waltham, MA). [α-$^{32}$P]-ATP (BLU003H250UC), [α-$^{32}$P]-CTP (BLU008H250UC), [α-$^{32}$P]-GTP (BLU006H250UC), [α-$^{32}$P]-UTP (BLU007H250UC), and S-[methyl-$^{14}$C]-Adenosyl-L-Methionine (NEC363010UC) were all obtained from PerkinElmer (Waltham, MA). All 5′-triphosphorylated RNAs were custom synthesized by ChemGenes Corporation (Wilmington, MA). Phenylmethylsulfonyl fluoride (PMSF; 97064-898) was obtained from VWR (Radnor, PA).

4–20% Mini-PROTEAN® TGX Stain-Free™ Protein Gels (4568096) were obtained from Bio-Rad. Uridine-5′-[(α,β)-imido]triphosphate (UMP-NPP; NU-930L) was obtained from Sapphire North America (Ann Arbor, MI).

Plasmids

SARS-CoV-2 nsp7, nsp8, nsp12, nsp13, nsp14, and nsp16 coding sequences (CDS) were codon-optimized for bacterial expression and synthesized as gBlocks (Integrative DNA Technologies, Coralville, IA). The CDS for nsp9 and nsp10 were amplified from mammalian expression vectors (a generous gift Nevan Krogen) [35]. The CDS were cloned into modified pET28a bacterial expression vectors containing N-terminal 6/8/10xHis tags followed by the yeast Sumo (smt3) CDS Amino acid mutations were introduced via QuikChange site-directed mutagenesis. Briefly, primers were designed using the Agilent QuikChange primer design program to generate the desired mutation and used in PCR reactions with PfuTurbo DNA polymerase. Reaction products were digested with Dpn1, transformed in DH5α cells and mutations were confirmed by Sanger sequencing.

For protein expression in *Escherichia coli* (*E. coli*), ppSumo-SARS-CoV-2 nsps and mutants were cloned into a BamH1 site at the 5′ end, which introduced a Ser residue following the diGly motif in smt3. To make native N-termini, the codon encoding the Ser was deleted via QuickChange mutagenesis. Thus, following cleavage with the ULP protease (after the diGly motif), the proteins contained native N-termini.

pGEX-2T-GST-eIF4E K119A [36] was obtained from Addgene (plasmid # 112818 )

### Protein purification

#### *Nsp5. nsp7, nsp8, nsp10, nsp14 and nsp16*

6xHis-Sumo-nsp5/7/8/10/14/16 and corresponding mutant plasmids (with native N-termini following the diGly motif in Sumo) were transformed into Rosetta (DE3) *E. coli* or LOBSTR-BL21(DE3)-RIL cells under 50 μg/ml kanamycin exposure. 5 ml LB Miller growth medium starter cultures containing 50 μg/ml kanamycin and 34 μg/ml chloramphenicol were grown for 2-4 h at 37°C and then transferred to growth medium containing the same antibiotics. Typically, 2 L were grown for each protein. Protein expression was induced at O.D. 0.7-1.0 by adding 0.4 mM IPTG and overnight incubation (16 hours) at 18°C. Cultures were centrifuged at 3,000 x g for 10 min and the bacterial pellet was resuspended in lysis buffer (50 mM Tris pH 8.0, 300 mM NaCl, 17.4 μg/ml PMSF, 15 mM imidazole pH 8.0 and 5 mM β-ME) and lysed by sonication. Lysates were centrifuged for 30 min at 30,000-35,000 x g and the supernatants incubated with Ni-NTA resin for 1-2 h at 4°C. The Ni-NTA resin was washed with 50 mM Tris pH 8.0, 300 mM NaCl, 30 mM imidazole pH 8.0,1 mM DTT and the protein was eluted in 50 mM Tris pH 8.0, 300 mM NaCl, 300 mM imidazole pH 8.0 and 1 mM DTT. The eluted proteins were incubated with 5 μg/ml Ulp1 protease overnight at 4°C. Nsp7 was separated from 6xHis-Sumo by anion exchange (Capto HiRes Q 5/50 column (Cytiva) equilibrated in 50 mM Tris 8.0, 50 mM NaCl, 1 mM DTT, eluted with 0-50% gradient of buffer containing 1 M NaCl). 6xHis-SUMO-nsp8 was treated with Ulp1 on the Ni-NTA resin, to separate nsp8 and 6xHis-SUMO in buffer with no imidazole. Proteins were further purified by size exclusion chromatography using Superdex 200 10/300 increase, Superdex 200 16/600, Superdex 75 10/300 increase or Superdex 75 16/600 columns in 50 mM Tris pH 7.5-8.0, 150-300 mM NaCl, 1 mM DTT, depending on yield and size. Fractions containing proteins of interest were pooled, concentrated in an Amicon Utra-15 with a 3-50 kDa pore size centrifugal filters. The nsp16 protein was incubated with Ni-NTA resin post SEC to remove 6His-SUMO. Purified proteins were aliquoted and stored at -80°C.

*Nsp9*

6xHis-Sumo-nsp9 and respective mutant plasmids (with native N-termini following the diGly motif in Sumo) were transformed into Rosetta (DE3) *E. coli* cells under 50 μg/ml kanamycin exposure. 5 ml LB Miller growth medium starter cultures containing 50 μg/ml kanamycin and 34 μg/ml chloramphenicol were grown for 2-4 h at 37°C and then transferred to Terrific Broth (TB) growth medium containing the same antibiotics and several drops of Antifoam B emulsion (Sigma, A5757). Protein expression was induced at O.D 1.2 by adding 0.4 mM IPTG and overnight incubation (16 hours) at 18°C. Cultures were centrifuged at 3,000 x g for 10 min and the bacterial pellet was resuspended in lysis buffer (50 mM Tris pH 8.0, 300 mM NaCl, 10% glycerol 17.4 μg/ml PMSF, 15 mM imidazole pH 8.0 and 5 mM β-ME) and lysed by sonication. Lysates were centrifuged for 30 min at 30,000 x g and the supernatants incubated in Ni-NTA resin for 1-2 h at 4°C. The Ni-NTA resin was washed with 50 mM Tris pH 8.0, 300 mM NaCl, 10% glycerol, 30 mM imidazole pH 8.0, 1 mM DTT and each protein was eluted in 50 mM Tris pH 8.0, 300 mM NaCl, 10% glycerol, 300 mM imidazole pH 8.0 and 1 mM DTT. The eluted protein was incubated with 5 μg/ml Ulp1 overnight at 4°C. Proteins were diluted with 50 mM Tris pH 8.0, 10% glycerol, 1 mM DTT buffer to lower the NaCl concentration to 30 mM and subsequently ran through a Hi-Trap CaptoQ column where the flowthrough contained purified nsp9. NaCl was added to each protein to a final concentration of 150 mM, concentrated in an Amicon Ultra-15 with a 10k MWCO, aliquoted, and stored at -80°C.

*Nsp12*

8xHis or 10xHis-Sumo-nsp12 and respective mutant plasmids (with native N-termini following the diGly motif in Sumo) were transformed into LOBSTR-BL21(DE3)-RIL *E. coli* cells under 50 μg/ml kanamycin exposure. 5 ml LB Miller growth medium starter cultures containing 50 μg/ml kanamycin and 34 μg/ml chloramphenicol were grown for 2-4 h at 37°C and then transferred to

1L growth medium containing the same antibiotics. Protein expression was induced at O.D. 0.8-1.2 by adding 0.4 mM IPTG and overnight incubation (16 hours) at 18°C. Cultures were centrifuged at 3,000-3.500 x g for 10 min and the bacterial pellet was resuspended in lysis buffer (50 mM Tris pH 8.0, 300 mM NaCl, 10% glycerol, 17.4 µg/ml PMSF, 15 mM imidazole pH 8.0 and 5 mM β-ME) and lysed by sonication. Lysates were centrifuged for 30 min at 30,000-35,000 x g and the supernatants incubated in Ni-NTA resin for 1-2 h at 4°C. The Ni-NTA resin was washed with high salt buffer: 50 mM Tris pH 8.0, 1 M NaCl, 10% glycerol, 30 mM Imidazole and 5 mM β-ME, followed by a high imidazole wash: 50 mM Tris pH 8.0, 300 mM NaCl, 10% glycerol, 75 mM Imidazole and 5 mM β-ME, and the protein was eluted in 50 mM Tris pH 8.0, 300 mM NaCl, 10% glycerol, 300 mM imidazole pH 8.0 and 1 mM DTT. The eluted proteins were incubated with 5 µg/ml Ulp1 overnight at 4°C. Proteins were further purified by size exclusion chromatography using a Superdex 200 10/300 increase column, or Superdex 200 16/600 in 50 mM Tris pH 8.0, 150-300 mM NaCl, 1 mM DTT. Fractions containing nsp12 were pooled, concentrated in an Amicon Ultra-15 with a 30-50k MWCO centrifugal filter, aliquoted, and stored at -80°C.

*Nsp13*

6xHis-Sumo-nsp13 (used in **Extended Data Fig. 12**), or 10xHis-Sumo-nsp13 (used for [$\alpha^{32}$P]-GTP conversion into GDP) and respective mutant plasmids (with native N-termini following the diGly motif in Sumo) were transformed into Rosetta (DE3) *E. coli* cells under 50 µg/ml kanamycin exposure. 5 ml LB Miller growth medium starter cultures containing 50 µg/ml kanamycin and 34 µg/ml chloramphenicol were grown for 2-4 h at 37°C and then transferred to 1 L of growth medium containing the same antibiotics. Protein expression was induced at O.D. 1.0 by adding 0.4 mM IPTG and overnight incubation (16 hours) at 18°C. Cultures were centrifuged at 3,000-3.500 x g for 10 min and the bacterial pellet was resuspended in lysis buffer (50 mM Tris pH 8.0, 300 mM NaCl, 10% glycerol, 17.4 µg/ml PMSF, 15 mM imidazole pH 8.0 and 5 mM β-ME) and lysed by sonication. Lysates were centrifuged for 30 min at 30,000-35,000 x g and the supernatants incubated in Ni-NTA resin for 1-2 h at 4°C. Ni-NTA resin for 6xHis-Sumo-nsp13 was washed with 50 mM Tris pH 8.0, 300 mM NaCl, 10% glycerol, 30 mM imidazole pH 8.0 and 1 mM DTT. 10xHis-Sumo-nsp13 with high salt buffer: 50 mM Tris pH 8.0, 1 M NaCl, 10% glycerol, 30 mM Imidazole and 5 mM β-ME, followed by a high imidazole wash: 50 mM Tris pH 8.0, 300 mM

NaCl, 10% glycerol, 75 mM imidazole and 5 mM β-ME, and the protein was eluted in 50 mM Tris pH 8.0, 300 mM NaCl, 10% glycerol, 300 mM imidazole pH 8.0 and 1 mM DTT. The eluted protein was incubated with 5 μg/ml Ulp1 overnight at 4°C. Proteins were buffer-exchanged or dialysed into a buffer containing 50 mM Bis-Tris pH 6.0, 30 mM NaCl, 10% glycerol and 1 mM DTT followed by ion-exchange chromatography in a 5/50 MonoS column. Fractions containing nsp13 were pooled and further purified by size exclusion chromatography using a Superdex 200 10/300 increase or Superdex 200 16/600 column in 50 mM Tris pH 8.0, 150 mM NaCl, 10% glycerol (only in 6xHis), 1 mM DTT. Fractions containing nsp13 were pooled, concentrated in an Amicon Ultra-15 with a 30-50k MWCO, aliquoted, and stored at -80°C.

*eIF4E*

For production of GST-eIF4E K119A, LOBSTR-BL21(DE3)-RIL cells were transformed with pGEX-2T-GST-eIF4E K119A [36] and were grown in LB supplemented with 100 μg/L Ampicillin, 34 μg/L chloramphenicol. Protein expression was induced at O.D. 1.0 by adding 0.4 mM IPTG and overnight incubation (16 hours) at 18 °C. Cultures were centrifuged at 3,000-3.500 x g for 10 min and the bacterial pellet was resuspended in lysis buffer (50 mM Tris pH 8.0, 300 mM NaCl, 17.4 μg/ml PMSF, and 5 mM β-ME) and lysed by sonication. Lysates were centrifuged for 30 min at 35,000 x g and the supernatants incubated with Pierce Glutathione resin for 1-2 h at 4 °C. The resin was washed with lysis buffer, and the GST-eIF4E K119A eluted with 50 mM Tris-HCl pH 8.0, 300 mM NaCl, 50 mM glutathione, 1 mM DTT. The protein was purified over Size Exclusion Chromatography on Superdex 200 16/600 in 50mM Tris-HCl pH 8.0, 300 mM NaCl, 1 mM DTT, concentrated, and stored as above.

*Ipp1*

For production of yeast inorganic pyrophosphatase (ipp1), The S. cerevisiae ipp1 CDS was cloned into pProEx2 containing a N-terminal 6xHis-TEV linker and was transformed into Rosetta *E. coli* cells under 100 μg/ml ampicillin exposure. 5 ml LB Miller growth medium starter cultures containing 100 μg/ml ampicillin were grown for 2-4 h at 37°C and then transferred to 1L growth medium containing the same antibiotics. Protein expression was induced at O.D. 0.8-1.2 by adding 0.4 mM IPTG and overnight incubation (16 hours) at 18°C. Cultures were centrifuged at 3,000-3.500 x g for 10 min and the bacterial pellet was resuspended in lysis buffer (50 mM Tris pH 8.0, 300 mM NaCl, 17.4 μg/ml PMSF, 15 mM imidazole pH 8.0 and 5 mM β-ME) and lysed by sonication. Lysates were centrifuged for 30 min at 35,000 x g and the supernatants incubated in Ni-NTA resin for 1 h at 4°C. Ni-NTA resin was washed with high salt buffer: 50 mM Tris pH 8.0, 1 M NaCl, 30 mM imidazole and 5 mM β-ME, and was eluted in 50 mM Tris pH 8.0, 50 mM NaCl, 300 mM imidazole pH 8.0 and 1 mM DTT. The eluted protein was loaded onto a Capto HiRes Q 5/50 column (Cytiva) equilibrated in 50 mM Tris 8.0, 50 mM NaCl, 1 mM DTT, eluted with 0-50% gradient of buffer containing 1 M NaCl. Protein was further purified by size exclusion chromatography using a Superdex 200 16/600 in 25 mM Tris pH 7.5, 50 mM NaCl, 2 mM DTT. Fractions containing YIPP were pooled, concentrated, and stored as above.

*Nsp8-nsp9 fusion*

The 6xHis-Sumo-nsp8-nsp9 plasmid and mutants (N1A, N1D and N2A) were transformed into Rosetta (DE3) *E. coli.* Cells were grown in Terrific broth media in the presence of 50 µg/ml kanamycin and 25 µg/ml chloramphenicol to OD 1.0 and induced with 0.4 mM IPTG for 16 hours at 18 °C. Cultures were centrifuged at 3500 x g for 15 minutes, and the pellets resuspended in lysis buffer (50 mM Tris, pH 8.0; 500 mM NaCl; 25 mM imidazole; 10% glycerol) in the presence of 1 mM PMSF. Cells were lysed by sonication and lysates cleared by centrifugation at 25000 x g for 1 hour. The lysate was passed over Ni-NTA beads, which were washed with lysis buffer. Protein samples were eluted with elution buffer (50 mM Tris, pH 8.0; 300 mM NaCl; 300 mM imidazole; 5% glycerol) and cleaved overnight at 4 °C with Ulp Sumo protease. Protein samples were further purified into cleavage assay buffer (50 mM Tris, pH 7.4; 150 mM NaCl; 5% glycerol) by size exclusion chromatography using a Superdex 75 Increase 10/300 GL column.

<u>Guanylyltransferase (GTase) activity assays</u>

GTase activity assays were performed as described in Yan et al. [21]. Reactions were assembled in 20 µL containing 50 mM Tris pH 8.0, 5 mM KCl, 1 mM DTT, 0.005 U/ml inorganic pyrophosphatase, 10 uM 5′-pppACCCCCCCCCCCCCCCCCCC-3’ (5′-pppRNA$^{A19C}$), 1.25 mM RNAse inhibitor, and where indicated, 0.5 µM nsp12, nsp12$^{D218A}$, nsp13, nsp13$^{K288A}$ or 1 U/ml of vaccinia capping enzyme (VCE). Reactions were started with 1 mM $MgCl_2$, 100 µM [α-$^{32}$P] GTP, (specific radioactivity = 1000 cpm/pmol) and incubated for 1 hr at 37°C. Half of the reaction was stopped by the addition of 0.8 U/ml proteinase K and incubated for 30 min at 37°C prior to the addition of 2X RNA loading dye (Novex) and incubated for 3 mins at 95°C. Reaction products were resolved in a 15% TBE-Urea PAGE gel. The gel was then stained with toluidine blue O and the $^{32}P$ signal detected via autoradiography.

The other half of the GTase reactions were treated with 10 U/ml P1 nuclease for 1 h at 37°C. Reactions were then split in half again with one half treated with 1 U/ml Quick CIP for 30 min at 37°C. Reactions were spotted on a PEI cellulose thin-layer chromatography (TLC) plate and developed in a 0.4 M ammonium sulfate $(NH_4)_2SO_4$ solvent system. The plate was dried and the $^{32}P$ signal was detected via autoradiography.

NMPylation assays

NMPylation reactions were carried out in 20 µL containing 50 mM Tris (pH 7.5), 5 mM KCl, 1 mM DTT, 16 µM nsp7, nsp8 or nsp9 (and mutants) and 4.8 nM nsp12 (and mutants). Reactions were started with 1 mM $MgCl_2$ or $MnCl_2$, 200 µM [α-$^{32}P$] ATP, [α-$^{32}P$] UTP, [α-$^{32}P$] GTP, or [α-$^{32}P$] CTP (specific radioactivity = 1000 cpm/pmol). The reactions were incubated at 37ºC for 5 minutes and stopped by adding 2 µL of 500 mM EDTA, followed by addition of 5X SDS-PAGE sample buffer with 10% β-ME and incubated for 3 minutes at 95°C. Reaction products were resolved by SDS-PAGE on a 4-20% gradient gel and visualized by staining with Coomassie Brilliant Blue. The $^{32}P$ signal was detected via autoradiography and scintillation counting.

Nsp9 NMPylation kinetics

NMPylation reactions were carried out in a 20 µL reaction containing 50 mM Tris (pH 7.5), 5 mM KCl, 1 mM DTT, 16 µM nsp9, and 4.8 nM nsp12. Reactions were started by adding $MnCl_2$ and [α-$^{32}P$] ATP, CTP, GTP, or UTP as indicated. The final concentration in the reaction was 0.5 to 200 µM (specific radioactivity = ~5000 cpm/pmol) of the indicated nucleotide triphosphate and 1 mM $MnCl_2$. The reactions were incubated at 37ºC for 5 minutes and stopped by adding 2 µL of 500 mM EDTA, followed by addition of 5X SDS-PAGE sample buffer + β-ME and boiling for 2-5 minutes. Reaction products were resolved by SDS-PAGE on a 4-20% gradient gel and visualized by staining with Coomassie Brilliant Blue. Incorporation of $^{32}P$ was quantified by excising the nsp9 bands from the gel and scintillation counting. Background radioactivity was subtracted from each measurement. Rate measurements were fit to Michaelis-Menten kinetic models and $K_m$ and $V_{max}$ for substrates were calculated by nonlinear regression using Prism 9.3.0 for macOS (GraphPad Software, San Diego, California USA, www.graphpad.com).

NMR

For NMR studies, non-isotopically enriched AMPylated nsp9 was dissolved in 50mM Tris buffer at pH 7.5, 150mM NaCl, 1mM DTT and 10% $D_2O$ for spectrometer locking. The final protein concentration of this solution was 0.5mM. A total volume of 500uL was then used with a 5mm NMR tube to record all the spectra.

All NMR experiments were run on a Bruker Avance III spectrometer operating at 600MHz (1H) and equipped with a 5mm proton-optimized quadruple resonance cryogenic probe. The temperature of the sample was regulated at 308K throughout data collection.

A one-dimensional (1D) 31P spectrum was recorded with 8192 scans and a repetition delay of 1.5sec for a total collection time of 3.5 hours. The 31P spectral window and offset were set to 17ppm and 2.6ppm, respectively. Waltz16 decoupling was used on 1H during 31P acquisition.

To observe contacts between 31P and the nearest 1H nuclei, a two-dimensional (2D) 1H,31P-HSQC spectrum was recorded with 1024 and 22 complex points in the direct 1H and indirect 31P dimensions, respectively. The spectral window and offset were set to 16.7ppm and 4.7ppm for the 1H dimension and 3.4ppm and 2.6ppm for the 31P dimension, respectively. Each FID was accumulated with 1536 scans with a repetition delay of 1sec for a total recording time of approximately 21 hours.

A 2D 1H,31P-HSQC-TOCSY spectrum was recorded using similar spectral window and offset parameters for the 1H and 31P dimensions as the HSQC spectrum described above. A 60ms long 1H-1H TOCSY pulse train using a DIPSI-2 sequence and a field strength of 10KHz was tagged at the end of the HSQC sequence to observe signals from 1H nuclei that are further away from 31P. Given the lower sensitivity of this experiment, each FID was accumulated with 4096 scans and a repetition delay of 1sec was used for a total recording time of 2 days and 14 hours.

Using a similar pulse sequence, a 2D 1H,1H-HSQC-TOCSY spectrum was also recorded by evolving the indirect 1H dimension instead of 31P. The spectral window for the 1H indirect dimension was set to 4.2ppm, while the offset was maintained at 4.7ppm as for the direct 1H dimension. 40 complex points were recorded for then indirect 1H dimension, using 2048 accumulations for each FID and a repetition delay of 1sec for a total recording time of 2 days and 14 hours.

All 2D spectra were processed using NMRPipe [37] and analysed with NMRFAM-SPARKY [38].

Intact mass analysis

Protein samples were analysed by LC/MS, using a Sciex X500B Q-TOF mass spectrometer coupled to an Agilent 1290 Infinity II HPLC. Samples were injected onto a POROS R1 reverse-phase column (2.1 x 30 mm, 20 μm particle size, 4000 Å pore size) and desalted. The mobile phase flow rate was 300 μL/min and the gradient was as follows: 0-3 min: 0% B, 3-4 min: 0-15% B, 4-16 min: 15-55% B, 16-16.1 min: 55-80% B, 16.1-18 min: 80% B. The column was then re-equilibrated at initial conditions prior to the subsequent injection. Buffer A contained 0.1% formic acid in water and buffer B contained 0.1% formic acid in acetonitrile.

The mass spectrometer was controlled by Sciex OS v.1.6.1 using the following settings: Ion source gas 1 30 psi, ion source gas 2 30 psi, curtain gas 35, CAD gas 7, temperature 300 ºC, spray voltage 5500 V, declustering potential 80 V, collision energy 10 V. Data was acquired from 400-2000 Da with a 0.5 s accumulation time and 4 time bins summed. The acquired mass spectra for the proteins of interest were deconvoluted using BioPharmaView v. 3.0.1 software (Sciex) in order to obtain the molecular weights. The peak threshold was set to ≥ 5%, reconstruction processing was set to 20 iterations with a signal-to-noise threshold of ≥ 20 and a resolution of 2500.

RNAylation assays

RNAylation reactions were typically carried out in a 10 μL volume containing 50 mM Tris (pH 7.5), 5 mM KCl, 1 mM DTT, 0.4 μg yeast inorganic pyrophosphatase, 20 μM nsp9, and 2 μM nsp12. Reactions were started by adding $MnCl_2$ and 5′-pppRNA$^{LS10}$ to a final concentration of 1 mM and 100 μM, respectively. Reactions were incubated at 37ºC for the indicated time points and stopped by addition of 5X SDS-PAGE sample buffer + β-ME and boiling the samples for 5 minutes. Reaction products were resolved by SDS-PAGE on a 4-20% gradient gel and visualized by Coomassie staining.

For the time course comparing the RNAylation of nsp9 N1A and N2A mutants (i.e. **Fig. 2d**), reactions were performed as above, except with 2.4 μM nsp12. At each indicated time point, reactions were stopped by addition of 5X SDS-PAGE sample buffer + β-ME and boiling the samples for 5 minutes. Reaction products were resolved by SDS-PAGE on a 4-20% gradient gel and visualized with Coomassie staining.

For reactions testing RNA length specificity (i.e. **Fig. 2e**), 7.5 µL of a reaction master mix containing nsp9, nsp12, and yeast inorganic pyrophosphatase was added to 2.5 µL of start mix consisting of $MnCl_2$ and the indicated 5′-pppRNA. The final reaction conditions were as follows: 50 mM Tris pH 7.5, 5 mM KCl, 1 mM DTT, 0.4 µg yeast inorganic pyrophosphatase, 20 µM nsp9, 2 µM nsp12, 1 mM $MnCl_2$, and 100 µM of the indicated RNA. Reactions were incubated for 30 minutes at 37ºC, then stopped by addition of 5X SDS-PAGE sample buffer + β-ME and boiling the samples for 5 minutes. Reaction products were resolved by SDS-PAGE on a 4-20% gradient gel and visualized by Coomassie staining.

For RNAylation reactions comparing different RNA sequences (i.e. **Fig. 2f**) or nsp9 mutants (i.e. **Fig. 5h**), reactions were performed as above, except using 1 µM nsp12. Reactions were incubated for 5 minutes and stopped by addition of 5X SDS-PAGE sample buffer + β-ME and boiling the samples for 5 minutes. Reaction products were resolved by SDS-PAGE on a 4-20% gradient gel and visualized by Coomassie staining.

<u>Purification of nsp9-pRNA$^{LS10}$ species</u>

Purified native nsp9 (0.8 mg/mL, 65 µM) was incubated at room temperature overnight with 130 µM of 5′-pppRNA$^{LS10}$ and ~0.8 µM of nsp12 in presence of 0.05 mg/ml yeast inorganic pyrophosphatase and 1 mM $MnCl_2$, in the reaction buffer (50 mM Tris 7.5, 5 mM KCl, 1 mM DTT). The samples were clarified by centrifugation to remove any precipitate and applied directly onto a Capto HiRes Q 5/50 column (Cytiva) equilibrated in 50 mM Tris 8.0, 50 mM NaCl, 1 mM DTT. An elution gradient of 0-50% with 1 M NaCl was applied over 30 column volumes. Under these conditions, RNA and nsp9-pRNA$^{LS10}$ bound the column and unmodified nsp9 did not. nsp9-pRNA$^{LS10}$ and unreacted RNA$^{LS10}$ eluted as a peak doublet around 70 mS/cm. Fractions were pooled, and further purified over Superdex 75 increase 10/300 GL (50 mM Tris 8.0, 300 mM NaCl, 1 mM DTT), separating nsp9-pRNA$^{LS10}$ from unreacted RNA$^{LS10}$, and nsp12. nsp9-pRNA$^{LS10}$ was quantified by spectrophotometry with an estimated extinction coefficient of $\epsilon_{260}$=130,650 $M^{-1}cm^{-1}$. We also generated nsp9-pRNA$^{LS10}$ in the absence of inorganic pyrophosphatase. This nsp9-pRNA$^{LS10}$ was used in control reactions to test for $PP_i$ hydrolysis, necessary to ensure that PPi mediated deRNAylation reactions shown in **Fig. 3e** do not suffer from pyrophosphate hydrolysis. The results were like those presented in the Figure, thus confirming that there was no contaminating inorganic pyrophosphatase in the assays.

DeRNAylation of nsp9-pRNA$^{LS10}$

DeRNAylation reactions were typically performed in a 10 µL reaction volume consisting of 50 mM Tris pH 7.5, 5 mM KCl, 1 mM DTT, 20 µM nsp9-pRNA$^{LS10}$, 1 µM nsp12, 1 mM $MgCl_2$, and 500 µM GDP. Reactions were started by adding $MgCl_2$/GDP and incubated at 37ºC for 5-60 minutes as indicated. Reactions were stopped by addition of 5X SDS-PAGE sample buffer + β-ME and boiling the samples for 5 minutes. Reaction products were resolved by SDS-PAGE on a 4-20% gradient gel and visualized with Coomassie staining.

For deRNAylation reactions comparing various nucleotide triphosphates (NTP) and nucleotide diphosphates (NDP), reactions were performed in 10 µL volume consisting of 50 mM Tris pH 7.5, 5 mM KCl, 1 mM DTT, 20 µM nsp9-pRNA$^{LS10}$, 500 nM nsp12, 1 mM $MgCl_2$, and 500 µM of the indicated NTP or NDP. Reactions were incubated for 5 minutes at 37ºC and stopped by addition of 5X SDS-PAGE sample buffer with β-ME and boiling for 5 minutes. Reaction products were resolved by SDS-PAGE on a 4-20% gradient gel and visualized with Coomassie staining.

Generation of [α-$^{32}$P]-GDP using nsp13

To generate [α-$^{32}$P]-GDP from [α-$^{32}$P]-GTP, 0.3-1 mM of [α-$^{32}$P]-GTP (specific activity ~2000 cpm/pmol) was incubated with 0.5-1 mg/mL nsp13 (and in some cases yeast cet1 NTPase) in 20 µL reaction buffer (depending on amount needed) consisting of 50 mM Tris (pH 7.5), 5 mM KCl, 1 mM DTT, 2 mM $MgCl_2$. Reactions were started by addition of enzyme and allowed to proceed for 30 minutes at 37ºC. Following the 30-minute incubation, reactions were boiled at 95ºC for 5 minutes to inactivate nsp13 or cet1.

Generation of radiolabelled GpppA-RNA$^{LS10}$ from nsp9-pRNA$^{LS10}$ and [α-$^{32}$P]-GDP

Reactions were performed in a 10 µL volume containing 50 mM Tris pH 7.5, 5 mM KCl, 1 mM DTT, 15 µM nsp9-pRNA$^{LS10}$, 377 nM nsp12, 1 mM $MgCl_2$, and 500 µM [α-$^{32}$P]-GDP (specific radioactivity = ~2000 cpm/pmol). Reactions were started by addition of [α-$^{32}$P]-GDP/$MgCl_2$ mixture (generated as described above) and incubated for 30 minutes at 37ºC. As a control, VCE was used but with [α-$^{32}$P]-GTP. VCE assays were generally performed as described in the NEB Capping Protocol (M2080) with the following modifications: the reaction contained 20 µM of 5'-pppRNA$^{LS10}$, 500 µM [α-$^{32}$P]-GTP (specific activity ~2000cpm/pmol), and did not contain SAM.

Reactions were stopped by the addition of 2X TBE-Urea sample buffer, boiled for 5 minutes, and resolved by UREA-PAGE (20%).

GDP inhibition of RNAylation (one pot capping assays)

Reactions were performed in 50 mM Tris pH 7.5, 5 mM KCl, 1 mM DTT, 1 mM $MgCl_2$, 1 mM $MnCl_2$, and contained 20 µM nsp9, 2 µM nsp12, and 100 µM 5'-pppRNA$^{LS10}$ in 20 µL volume. The [α-$^{32}$P]-GDP was prepared as described earlier (using 400 µM GTP, [α-$^{32}$P]-GTP at specific activity ~1,500 cpm/pmol) and diluted to final reaction concentrations in the range of 6.25-100 µM. Reactions were started by the addition of nsp12. GDP was added either before the addition of nsp12 (t=0), or after 30 minutes of preincubation. After an additional 30 minutes, the reactions were split in half and stopped by the addition of 5x SDS-PAGE or 2x Formamide loading dyes and the products were analysed by 4-20% gradient SDS-PAGE gel, or 15% 19:1 TBE UREA-PAGE gel, respectively.

LC-MS/MS analysis of GpppA

De-RNAylation reactions (in triplicate) were performed in 20 µL of buffer solution containing 50 mM Tris pH 7.5, 5 mM KCl, 20 µM nsp9-pRNA$^{LS10}$, 2 µM wild-type nsp12 or the D218A mutant, 1 mM $MgCl_2$, and 100 µM GDP. Reactions were started by adding $MgCl_2$ and GDP and allowed to proceed for 1 hour at 37ºC. After the 1-hour incubation, the reactions were supplemented with 2 µL of 10X P1 buffer and 1 µL of Nuclease P1 enzyme and allowed to proceed for an additional 30 minutes at 37ºC. Reactions were stopped by boiling for 5 minutes and submitted for LC-MS/MS analysis.

For standards, 20 µL of blank reaction buffer (50 mM Tris pH 7.5, 5 mM KCl, 1 mM DTT) plus 40 µL blank reaction buffer containing 0.33 µM (final) m7GpppA (m7G(5′)ppp(5′)A RNA Cap Structure Analog, New England Biolabs, S1405S) as an internal standard (IS) was spiked with varying concentrations of GpppA (New England Biolabs, S1406L). Reaction samples (20 µL) were diluted with blank reaction buffer, containing 0.33 µM (final) m7GpppA IS, at 1:2 to a total volume of 60 µL. Standards and samples were mixed with 60 µL of 100% methanol, vortexed, and then spun for 5 min at 16,100 x g. Supernatant was removed and analysed by LC-MS/MS using a Sciex (Framingham, MA) QTRAP® 6500+ mass spectrometer coupled to a Shimadzu (Columbia, MD) Nexera X2 LC. GpppA was detected with the mass spectrometer in positive MRM (multiple reaction monitoring) mode by following the precursor to fragment ion transition

772.9 → 604.0. A Thermo Scientific BioBasic AX column (2.1 x 50 mm, 5 micron packing) was used for chromatography with the following conditions: Buffer A: 8:2 $dH_2O$:Acetonitrile + 10 mM ammonium acetate, pH 6, Buffer B: 7:3 $dH_2O$:Acetonitrile + 1 mM ammonium acetate, pH 10.5, 0.5 mL/min flow rate, 0-1 min 0%B, 1-2.5 min gradient to 35%B, 2.5-5 min 35%B, 5-7 min gradient to 65%B, 7-10 min 65%B, 10-10.5 min gradient to 100%B, 10.5-15 min 100%B, 15-15.5 min gradient to 0%B, 15.5-20.5 min 0%B. m7GpppA (transition 787.1 → 508.0) was used as an internal standard. Peak areas were determined and data were further analysed using the Sciex Analyst 1.7.2 software package. Back-calculation of standard curve samples were accurate to within 15% for 100% of these samples at concentrations ranging from 0.001 µM to 10 µM. A limit of detection (LOD) was defined as a level three times that observed in blank reaction buffer and the limit of quantitation (LOQ) as the lowest point on the standard curve that gave an analyte signal above the LOD and within 20% of nominal upon back-calculation. The LOQ for GpppA was 0.005 µM.

Methyltransferase assays

In 10 µL reactions, 40 µM nsp9-$pRNA^{LS10}$ was incubated with 2 µM nsp12 in presence of 1 mM $MgCl_2$ and 100 µM $[^{32}P]$GDP (generated as above) for 60 min at 37 °C. Reactions were filled to 15 µL with nsp14 and SAM (final 0.05 mg/mL and 100 uM respectively), and incubated for another 30 minutes. Treatment with nsp10/16 can be done concurrently with nsp14, however nsp10/14 complex partially processes $RNA^{LS10}$, resulting in a mobility shift [39]. Thus, prior to addition of nsp10 and nsp16, nsp14 exonuclease activity was removed by heat inactivation (5 minutes at 95 °C). For 2'-O methylation, reactions were supplemented with nsp10/16 and fresh SAM to final concentrations of 0.05 mg/mL nsp10, 0.05 mg/mL nsp16, 100 µM SAM in final 20 µL volume. Vaccinia reactions were conducted as per manufacturer's instructions. Reactions were stopped by adding 2x formamide loading dye and were separated on 20% TBE-UREA polyacrylamide gels (19:1). Radioactivity was visualized by autoradiography and RNA by toluidine staining. For TLC analysis, bands with detectible $^{32}P$ signal were excised, fragmented, and incubated overnight at 55°C in elution buffer (1 M Ammonium acetate, 0.2% SDS, 20 mM EDTA), rotating top-over-bottom. Solutions were filtered using 0.22 µm centrifugal filters, supplemented with 23 ug Glyco Blue co-precipitant (Invitrogen) and precipitated for 1 hr at -20°C by addition of isopropyl alcohol to a final concentration of 60%. The pellets were washed once with 70% EtOH and reconstituted in 10 µL of P1 buffer with P1 nuclease (NEB). After 30 minutes at 37°C, reactions were supplemented with Quick CIP (NEB) and rCut Smart buffer, to a final volume of 12 uL. After 30 min of further incubation, reactions were spotted onto PEI-Cellulose F TLC plates and resolved in 0.4 M Ammonium Sulfate mobile phase. Beforehand, TLC plates were prepared by development in water, removing yellow discoloration. The $^{32}$P signal was detected by autoradiography, and compared with cold standards of GTP, GDP, GpppA and $^{m7}$GpppA detected by absorption of plate fluorescence, excitable with a UV lamp λ=265 nm. The position of $^{m7}$GpppA$_{2'\text{-OMe}}$ was determined from the Vaccinia capping enzyme and Vaccinia 2'-O-Methyltransferase control reaction.

Reactions with $^{14}$C-labelled SAM were conducted as above, with two differences: cold GDP was used at 100 μM (Millipore Sigma, G7127), and 55 μM [$^{14}$C ]SAM (Perkin Elmer), SAM was used at the supplied radioactivity of 52.6 mCi/mmol (~117 cpm/pmol), with no further dilution using cold SAM.

GST-eIF4E pulldown of $^{7Me}$GpppA-RNA

Capping reactions were set up in 20 μL and contained 2 μM nsp12, 0.04 mg/ml nsp14 WT or D331A, 30 μM nsp9-pRNA$^{LS10}$, 100 μM [α$^{32}$P]GDP (specific radioactivity = 1,000 cpm,/pmol), 100 μM SAM, 2 mM $MgCl_2$. The reaction buffer was 50 mM Tris 8.0, 5 mM KCl, 1 mM DTT. Vaccinia capping enzyme controls were performed according to manufacturer's instructions, with the same nucleotide and SAM concentrations as nsp12 reactions. After 90 minutes of incubation at 37 °C, 15.6 ug of GST-eIF4E K119A was added, along with 15 μL of Glutathione resin (Pierce, Thermo Scientific). Reactions were filled to 700 μL with 50 mM Tris 8.0, 150 mM NaCl, 1 mM DTT, and nutated for 1 hour. The resin was washed 3x with 500 μL of 50 mM Tris 8.0, 150 mM NaCl, 1 mM DTT, and radioactive signal was quantified by scintillation counting.

Cryo-EM grid preparation

To form nsp12/7/8 core complex (RTC), native nsp12, nsp7 and nsp8 were incubated in 1:2:4 molar ratio and run over Superdex 200 increase 10/300 GL to separate unassociated monomers. The purified complex was concentrated using spin concentrators (Amicon 10k MWCO, Sigma-Millipore), and quantified by spectrophotometry. A 3x molar excess of nsp9 over RTC was added, followed by 0.05 mM final DDM detergent immediately prior to freezing. Final concentration of nsp12/7/8 was 2 mg/mL. Buffer contained 50 mM Tris 7.5, 150 mM NaCl, 1 mM DTT, 2 mM

$MnCl_2$, 1 mM UMP-NPP. Copper Quantifoil 1.2/1.3 mesh 300 grids were used to freeze 3.5 μL of sample at 100% relative humidity using Vitrobot mk. IV (Thermofisher).

<u>Cryo-EM data collection</u>

Prior to data collection, sample grids were screened on a Talos Artica microscope at the Cryo Electron Microscopy Facility (CEMF) at UT Southwestern. Cryo-EM data of NSP12/7/8/9 complex were collected on a Titan Krios microscope at Cryo-Electron Microscopy Facility (CEMF) at UT Southwestern Medical Center, with the post-column energy filter (Gatan) and a K3 direct detection camera (Gatan), using SerialEM [40]. 4,770 movies were acquired at a pixel size of 0.55 A in super-resolution counting mode, with an accumulated total dose of 54 e-/$Å^2$ over 50 frames. The defocus range of the images was set to be -1.0 to -2.5 μm.

<u>Image processing and 3D reconstruction</u>

Unless described otherwise, all datasets were processed with Relion [41]. Movies were aligned and summed using MotionCor2 [42], with a downsampled pixel size of 1.09 Å. The CTF parameters were calculated using Gctf [43], and images with estimated CTF max resolution better than 5 Å were selected for further processing. 4,196,086 particles were picked using crYOLO [44] from 4,757 images, and extracted with a re-scaled pixel size of 2.19 Å. 663,999 particles were selected and re-extracted after multiple rounds of 2D and 3D classifications in Relion with the original pixel size of 1.09 Å. An additional round of 3D classification was carried out, followed by particle reduction with a homemade script to remove particles from dominant orientations. The remaining 89,945 particles were subjected to 3D refinement, CTF refinement and particle polishing sequentially. A final round of 3D classification with a reference mask led to 39,985 particles, which were then imported into cryoSPARC [45] for one round of non-uniform refinement. The map resolution was reported at 3.18 Å from cryoSPARC with the gold standard FSC method.

<u>Nsp5 cleavage reactions</u>

Concentrated protein samples were diluted in cleavage buffer (50 mM Tris, pH 7.4; 150 mM NaCl; 5% glycerol) and each reaction was performed in a total volume of 10 μL. Initial experiments measured the nsp5 concentration dependence of the nsp8-nsp9 cleavage reaction. To measure time dependence, nsp5 (2.5 μM final concentration) was added to the nsp8-nsp9 fusion protein (12.5 μM final concentration). Reactions were incubated at 37 °C for varying amounts of time (0 to 80 minutes) and terminated by boiling the samples for five minutes in the presence of SDS-PAGE loading buffer. Reaction products were resolved on a 4-20% gradient tris-glycine gel and products were visualized by Coomassie staining.

Model building and refinement

Model was build using PDB 7CYQ as a template [21]. Model was manually rebuilt into the map using Coot [46], and refined using Phenix real space refinement [47]. Model validation was performed using MolProbity software [48].

Bioinformatics

A representative subset of NiRAN domain sequences, provided as a multiple alignment in the original NiRAN publication [5], was supplemented by additional sequences used in this study (SARS-CoV-2, OC43, 229E strains). The human SELO sequence was added according to a FATCAT structural alignment [49] between SARS-CoV-2 nsp12 and bacterial SelO (PDB identifiers 7cyq and 6eac). The alignment was visualized using the ESPript server [50].

SARS-CoV-2 infection experiments

*Plasmid Construction*

To generate recombinant SARS-CoV-2 expressing ZsGreen mutants, the infectious clone pCC1-4K-SARS-CoV-2-Wuhan-Hu-1-ZsGreen was used as the parental backbone [8]. To generate nsp9 (N1A, N1D, N2A) and nsp12 (K73A) mutants, mutations were introduced by overlap extension PCR. Briefly, 2 fragments for each mutant were created by PCR using Ex Taq DNA Polymerase (Takara). These fragments shared homology at the 3' end of Fragment 1 and 5' end of Fragment 2. The resulting 2 fragments were then used as a template for a third fragment to PCR a full-length amplicon containing the mutation flanked by Pac1 and MluI sites on the ends. Mutations were confirmed by DNA sequencing. To generate nsp12 mutants (D218A, D760A), a SARS-CoV-2 shuttle vector (ps1180.SARS-CoV-2-shuttle) was created using ps1180.delXhoISacII plasmid as the backbone. Using Gibson cloning, a restriction enzyme linker that contained unique restriction sites specific to the SARS-CoV-2 genome was inserted. Smaller fragments of the SARS-CoV-2 genome were digested from the pCC1-4K-SARS-CoV-2-Wuhan-Hu-1-ZsGreen plasmid and ligated into ps1180.SARS-Cov-2-shuttle plasmid to create 3 new plasmids (MluI/SacI fragment for D218A region; SacI/Bsu36I fragment for D760A region). gBlocks containing mutations were synthesized by IDT and introduced into the SARS-CoV-2 shuttle vector by Gibson Assembly following standard protocols. To reassemble the full length parental pCC1-4K-SARS-CoV-2-Wuhan-Hu-1-ZsGreen containing new mutants, pCC1-4K-SARS-CoV-2-Wuhan-Hu-1-ZsGreen was digested with PacI/MluI, MluI/SacI, and SacI/Bsu36I restriction enzymes. A roughly 28-35 kb fragment was purified for each digest. PCR Amplicons ($nsp9^{N1A}$, $nsp9^{N1D}$, $nsp9^{N2A}$, $nsp12^{K73A}$) were digested with PacI/MluI and 5.3 kb fragment was purified. The SARS-CoV-2 shuttle plasmids were digested as follows: $nsp12^{D218A}$ (MluI/SacI releasing 1.4 kb fragment); $nsp12^{D760A}$ (SacI/Bsu36I/PvuI releasing 3 kb fragments). All fragments were purified using QIAexII Gel Purification Kit following standard protocol (Qiagen). Fragments were ligated together at a 3:1 ratio overnight. Ligated DNA was precipitated using 7.5 M Ammonium acetate, Glycogen, and Isopropanol, followed by an ethanol wash. The DNA was electroporated into TransforMAX EPI300 Electro competent *E. coli* (Lucigen). An overlapping 8-fragment PCR strategy was used to verify individual colonies by colony PCR. Confirmed colonies were grown in 10 ml Tryptic Soy Broth (TSB; Sigma) containing 12.5 µg/ml Chloramphenicol for 6-8 hours, shaking at 37°C. A 10 ml culture was inoculated into 100 ml TSB/Chloramphenicol culture and incubated overnight, shaking at 37°C. Overnight culture was diluted 1:5 into fresh TSB/Chloramphenicol containing 0.1% Arabinose and incubated an additional 5 hrs. Bacteria was pelleted and DNA was isolated using a homemade midi prep protocol followed by Machery-Nagel NucleoBond Xtra Midi Kit (Fisher). Full-length infectious clone plasmid was confirmed by restriction digestion and 8-fragment PCR. Oligonucleotides used are shown in **Table S3**.

*Virus production*

To generate virus from DNA-based infectious clones, 3 µg of plasmid were transfected into 2 individual 6 wells of 400,000 BHK-21J cells using X-treme Gene9 Transfection Reagent (Sigma). Three days post transfection, the supernatant from 2 individual wells was combined and 3 ml was transferred to a T25 flask containing 1 x $10^6$ VeroE6-C1008-TMPRSS2 cells and 2 ml serum-free MEM. After 4 days, 250 µl supernatant was added to 750 µl TriReagent for RNA extraction and RT-qPCR. T25 flasks were fixed with 4% paraformaldehyde, imaged on Nikon Eclipse Ti and processed with ImageJ.

*RNA Extraction/RT-qPCR*

RNA was isolated using the Direct-zol RNA mini prep kit following manufacturer's instructions (ZymoResearch). A 20 µl reaction contained 5 µl RNA, 5 µl TaqMan Fast Virus 1-Step Master Mix, and 1.8 µl SARS-CoV-2 primer/probe set containing 6.7 µM each primer/1.7 µM probe (final concentration of primer/probe were 600 nM/150 nM probe). SARS-CoV-2 primers and probe were designed as recommended by the Center for Disease Control (https://www.cdc.gov/coronavirus/2019-ncov/lab/rt-pcr-panel-primer-probes.html). All oligonucleotides were synthesized by LGC Biosearch Technologies. RT was performed at 50°C for 5 minutes, followed by inactivation at 95°C for 2 minutes, and 40 cycles of PCR (95°C for 3 seconds, 60°C for 30 seconds) on a QuantStudio 3 (Applied Biosystems).

*Cells*

BHK-21J cells (a generous gift from C. Rice) were grown in MEM (Gibco) supplemented with 10% FBS and 1X NEAA. VeroE6-C1008 cells (ATCC) were transduced with lentiviral vector SCRBBL-TMPRSS2, selected and maintained in MEM supplemented with 10% FBS, 1X NEAA, and 8 µg/ml Blasticidin.
